# Ventral arkypallidal neurons modulate accumbal firing to promote reward consumption

**DOI:** 10.1101/2020.04.01.020099

**Authors:** Yvan M. Vachez, Jessica R. Tooley, Eric Casey, Tom Earnest, Kavitha Abiraman, Hannah Silberberg, Elizabeth Godynyuk, Olivia Uddin, Lauren Marconi, Claire Le Pichon, Meaghan C. Creed

**Author notes:** Equal contribution (Yvan M. Vachez, Jessica R. Tooley). Correspondence, @Meaghan_Creed.

## Abstract

A pause in firing of nucleus accumbens shell (NAcSh) neurons is critical for reward consumption; however, the substrate driving this pause is unknown. While ventral pallidal (VP) activity encodes reward value, the specific roles of VP subpopulations in computation and expression of this value are poorly understood. Here, we establish that inhibitory input from the VP is crucial for reward-related inhibition of NAc firing. A sparse, non-canonical subpopulation of VP neurons, the so-called “ventral arkypallidal (vArky)” neurons makes inhibitory synaptic contacts throughout the NAcSh, and drives inhibition of NAcSh neurons in vivo. Moreover, endogenous calcium activity of vArky neurons predicted subsequent reward consumption behavior, while optogenetically activating this pathway increased reward consumption by amplifying hedonic value of reward. Classically, the VP is considered downstream of the NAc; however, our results challenge this view and establish that vArky neurons in the VP promote reward consumption via potent modulation of NAcSh firing.

## INTRODUCTION

The ventral basal ganglia is a collection of interconnected subcortical nuclei which are critical for reward processing and generating goal-directed behaviors (Humphries & Prescott, 2010). The nucleus accumbens (NAc) and ventral pallidum (VP) are two key nuclei within this network, and are crucial for hedonic valuation and reward consumption (Castro et al., 2015). The NAc shell (NAcSh) is comprised of medium spiny neurons expressing the D1- or D2-dopamine receptor (D1-MSNs, D2-MSNs, respectively), which both densely innervate the VP (Creed et al., 2016; Kupchik et al., 2015). The VP is therefore considered the primary output nucleus of the NAcSh (Nauta et al., 1978; Williams et al., 1977). NAcSh neurons are inhibited during reward consumption, and this inhibition is both necessary and permissive for reward consumption (O’Connor et al., 2015). Moreover, the magnitude of this inhibition scales with reward value (Faure et al., 2008; Krause et al., 2010; Roitman et al., 2005; Zhang et al., 2003), and is hypothesized to produce motivational drive by disinhibition of downstream targets, including the VP (Heimer et al., 1991; Humphries & Prescott, 2010; Smith et al., 2011; Stratford, 2005). However, the neural substrate driving the maintenance of this pause in NAcSh firing is still poorly understood.

The unidirectional organization of information flow from NAc to VP has been called into question by neuroanatomical studies and recent functional evidence. Specifically, tract-tracing has demonstrated the existence of projections from the VP to the NAc in rodents (Brog et al., 1993; Churchill & Kalivas, 1994; Groenewegen et al., 1993) and primates (Spooren et al., 1996). Functionally, value-related neuronal firing is more widespread in the VP compared to NAc, while firing responses to reward-consumption and to cues that predict reward occur with shorter latency in the VP neurons relative to NAc neurons (Ambroggi et al., 2011; Ottenheimer et al., 2018; Richard et al., 2016). In the dorsal basal ganglia, projections from globus pallidus to striatum have been coined “arkypallidal”, due to their dense net-like projections impinging broadly through the dorsal striatum (Glajch et al., 2016; Mallet et al., 2012, 2016). We hypothesize that an analogous ventral arkypallidal pathway (vArky; from VP to NAcSh) is one neuronal substrate by which reward-related activity in the VP can drive reward-related inhibition in NAcSh firing, and could reconcile the different latencies with which reward-related signals arise in these two nuclei.

Here, we anatomically and functionally characterize the vArky pathway. These vArky neurons send dense inhibitory projections to NAcSh MSNs and interneurons, and this inhibitory drive contributes to reward-related neuronal responses in the NAcSh. The endogenous activity of vArky neurons predicts subsequent reward consumption, and promotes reward consumption by increasing hedonic value of reward. We thus describe the first distinct behavioral function of ventral arkypallidal neurons, while providing new insights into information flow in the ventral basal ganglia necessary for understanding reward processing.

## RESULTS

### NAcSh inhibition promotes reward consumption

Inhibition of the NAcSh during reward consumption has been consistently documented, although there is heterogeneity across paradigms regarding operant contingencies, method of responding or metabolic state (Cheer et al., 2005; Krause et al., 2010; Roitman et al., 2010; Taha & Fields, 2005, 2006). Here, in free-access paradigm, where freely moving, stated mice drink from a lickometer, 35.7% of recorded NAcSh neurons were inhibited during reward consumption (Fig1A-D). Also consistent with prior work, the response during reward approach were more heterogeneous, with 7.14% of neurons showing a decrease, 25.0% an increase and 67.86% no change (Fig S1A-D). Further, NAcSh inhibition is sufficient to drive reward consumption, since closed-loop optogenetic inhibition with Arch3.0 triggered by reward consumption increased the total reward consumption time, number of bouts and average bout length (Fig1E-H, S2A-D). NAc inhibition did not induce a place preference or induce locomotor effects (Fig S2E,F). These experiments establish that activation of the vArky pathway is sufficient to inhibit NAcSh neurons, and that inhibition of NAcSh neurons occurs during reward consumption. We hypothesized that the VP is an important exogenous source of inhibition to the NAcSh that could drive this pause in firing, given its established role in encoding motivation and reward value (Ledbetter et al., 2016; Ottenheimer et al., 2018; Tachibana & Hikosaka, 2012).

**Figure 1.**
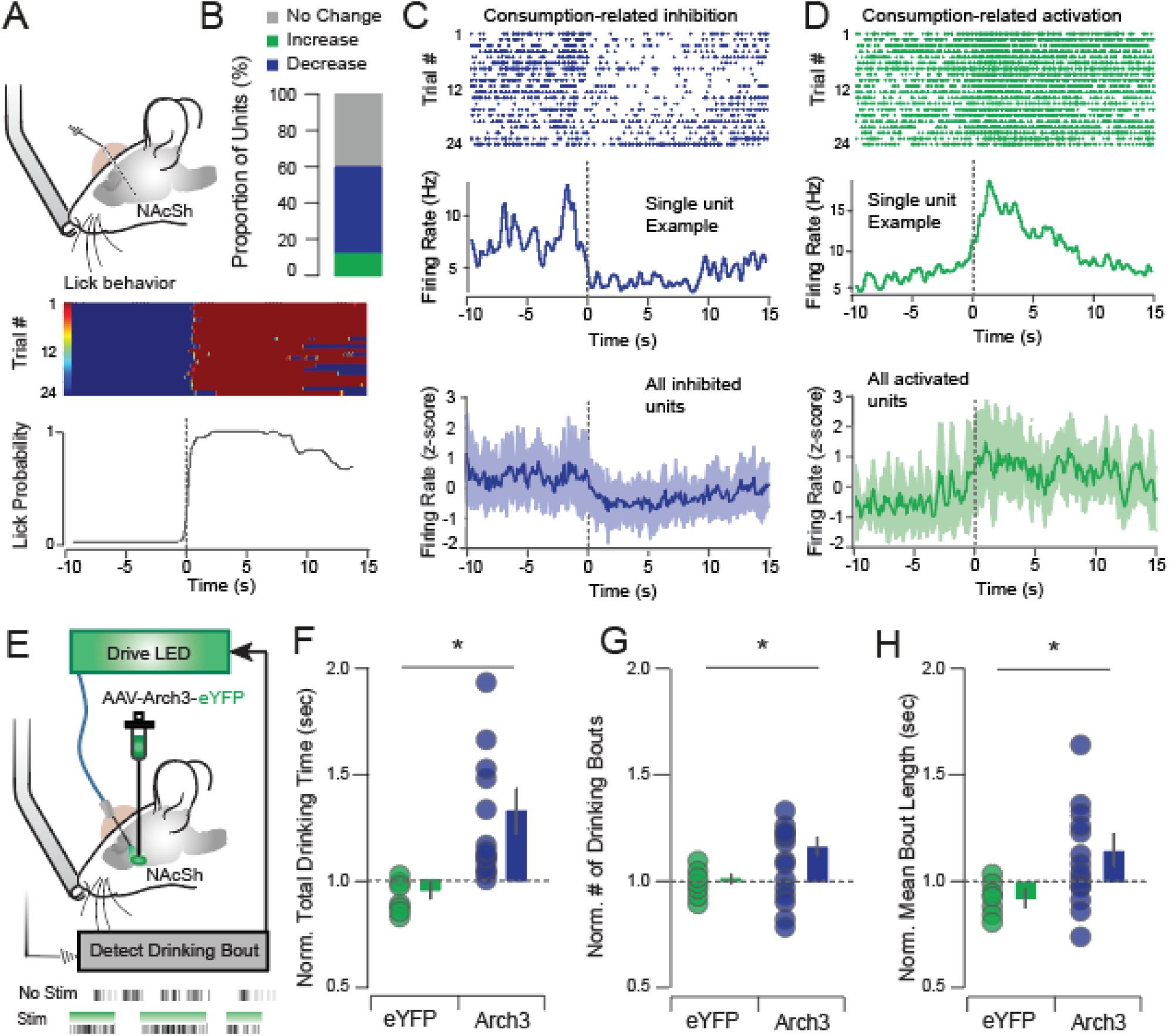
NAcSh inhibition is causally related to reward consumption. **(A)** In vivo recording of the NAcSh during free-access paradigm (n=32 multiunits, 6 mice). **(B)** Summary of drinking-related responses of NAcSh units; 35.71% decrease, 10.71% increase, 53.57% no change. **(C-D)** Top: examples of reward-inhibited and reward activated single units from the same mouse, aligned to drinking onset. Raster and PSTH of units inhibited (C) and activated (D) during reward consumption; bottom = mean ± SEM. **(E)** Closed-loop optogenetic inhibition of NAcSh neurons during reward consumption (n=12/15 eYFP/Arch3). **(F-H)** Optogenetic inhibition increased total drinking time (Arch: 1.29 ± 0.11, eYFP: 0.95 ± 0.03), number of drinking bouts (Arch: 1.01 ± 0.02, eYFP: 1.16 ± 0.06) and bout length (Arch: 1.12 ± 0.06, eYFP: 0.93 ± 0.02) relative to eYFP controls. * p < 0.05

### vArky neurons inhibit the NAcSh

To trace projections from the VP to NAcSh, we injected retrograde-AAV expressing Cre into the NAcSh, and cre-dependent ChR2-tagged to eYFP into the VP (Fig S3A). We confirmed that cell bodies in the subcommisural VP were labelled, and that eYFP+ terminals were abundant in the NAcSh (Fig S3B,C); fluorescence signal and length of labeled dendrites in the NAcSh was significantly greater than all other brain regions examined (Fig S3D,E, Vid1). We refer to these NAcSh-projecting VP neurons as ventral arkypallidal (vArky) neurons, given their dense, net-like morphology (Vid1), which is reminiscent of the arkypallidal projection from globus pallidus to striatum in the dorsal basal ganglia (Mallet et al., 2012, 2016).

We and others have recently shown that glutamatergic VP neurons constitute a distinct population of projection neurons (Faget et al., 2018; Tooley et al., 2018). To determine whether the vArky pathway is comprised of GABAergic or glutamatergic neurons, we injected retrobeads in the NAcSh and measured co-expression of VP retrobeads with SLC32A1 (encoding vesicular inhibitory amino acid transporter, VIAAT) and SLC17A6 (vesicular glutamate transporter 2, VGluT2) mRNA using fluorescent in situ hybridization (fISH, Fig 2A,B). We found 86.6% of retrobead-positive neurons co-localized with VIAAT consistent with early anatomical studies (Churchill & Kalivas, 1994); there was no co-localization between retrobeads and VGluT2 (Fig 2C).

**Figure 2.**
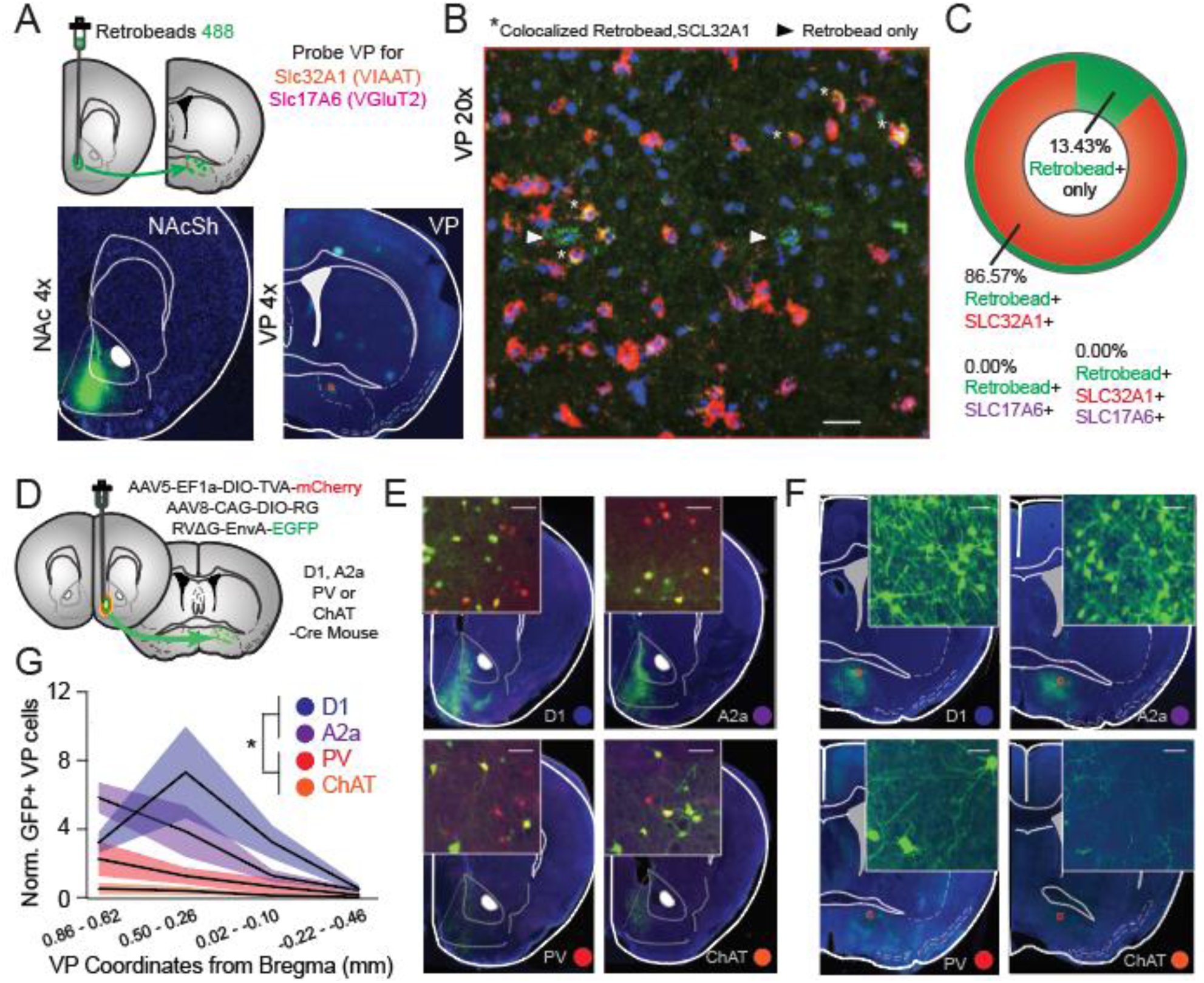
A subpopulation of VP neurons innervates projection and interneurons in the NAcSh. **(A)** Green retrobeads were injected in the NAcSh, VP sections were probed for VIAAT (555 nm, red) and VGluT2 (647 nm; magenta) using fISH. (**B)** Representative 20x confocal image showing green-labelled retrobeads in the VP co-localized with VIAAT. **(C**) Quantification of overlap between retrobeads and VIAAT (86.57% of retrobead+ neurons) and VGluT2 (0%, no co-localization). **(D)** Schematic of rabies tracing viral injection strategy. **(E)** Representative images of injection sites in NAcSh in D1-Cre (n = 5 mice), A2a-Cre (n = 5 mice), PV-Cre (n = 5 mice) and ChAT-Cre (n = 7) mice; inset = 20x confocal images showing starter cells (scale bar = 50 µM). **(F)** Representative images of EGFP-infected projection neurons in the VP; inset = 20x confocal images (scale bar = 50 µM). **(G)** Quantification of the number of EGFP+ VP cells, normalized to number of starter cells in the NAcSh is expressed across the anterior-posterior VP gradient revealed a significant effect of genotype, with D1- and A2a-Cre mice exhibiting denser labelling than PV-Cre or ChAT-Cre mice (F_15_ = 13.56, p<0.001, post-hoc t-test, D1-Cre = 3.67 ± 0.79, A2a-Cre = 3.50 ± 0.81, PV-Cre = 1.07 ± 0.37, ChAT-Cre = 0.25 ± 0.10 EGFP+ive cells/starter cell). *p<0.01.

The NAcSh is composed D1- and D2-MSNs, as well as interneurons. To determine the post-synaptic targets of vArky neurons within the NAcSh, we used complementary approaches of retrograde genetic tracing and anterograde circuit mapping. Retrograde tracing using G-deleted rabies virus seeding in the NAcSh (Fig 2D) revealed that normalized counts of EGFP-labelled cells in VP were significantly denser in D1-Cre and A2a-Cre mice relative to PV-Cre and ChAT-Cre mice (Fig 1E-G), reflecting increased monosynaptic inputs from VP onto MSNs relative to interneurons. Consistent with prior anatomical tracing, our data reveal that these neurons were concentrated around the anterior-medial, subcomissural VP, with only sparse labeling in caudal VP (Churchill & Kalivas, 1994; Fig 2G). We corroborated these results using functional circuit mapping, by injecting ChR2 into the VP, and recording light-evoked responses in NAcSh D1-MSNs, D2-MSNs, PVs or CINs using patch-clamp electrophysiology (Fig 3A,B and S4A,B). Corroborating our rabies tracing results, 84% of D1-MSNs and 79% of D2-MSNs received monosynaptic input from the VP, whereas only 30% of CINs and 28% of PVs received monosynaptic VP input (Fig 3C,D). There was no difference in connectivity rate (Fig 3D), amplitude or release probability (Fig 3E, S4C,D) as a function of location within the NAcSh. The latency to current peak and the insensitivity of this latency to tetrodotoxin confirm that these currents are monosynaptic (Fig 3F,G). Consistent with our fISH results, light-evoked currents were blocked by picrotoxin, but not by APV/NBQX, confirming their exclusively GABAergic nature (Fig 3H,I).

**Figure 3.**
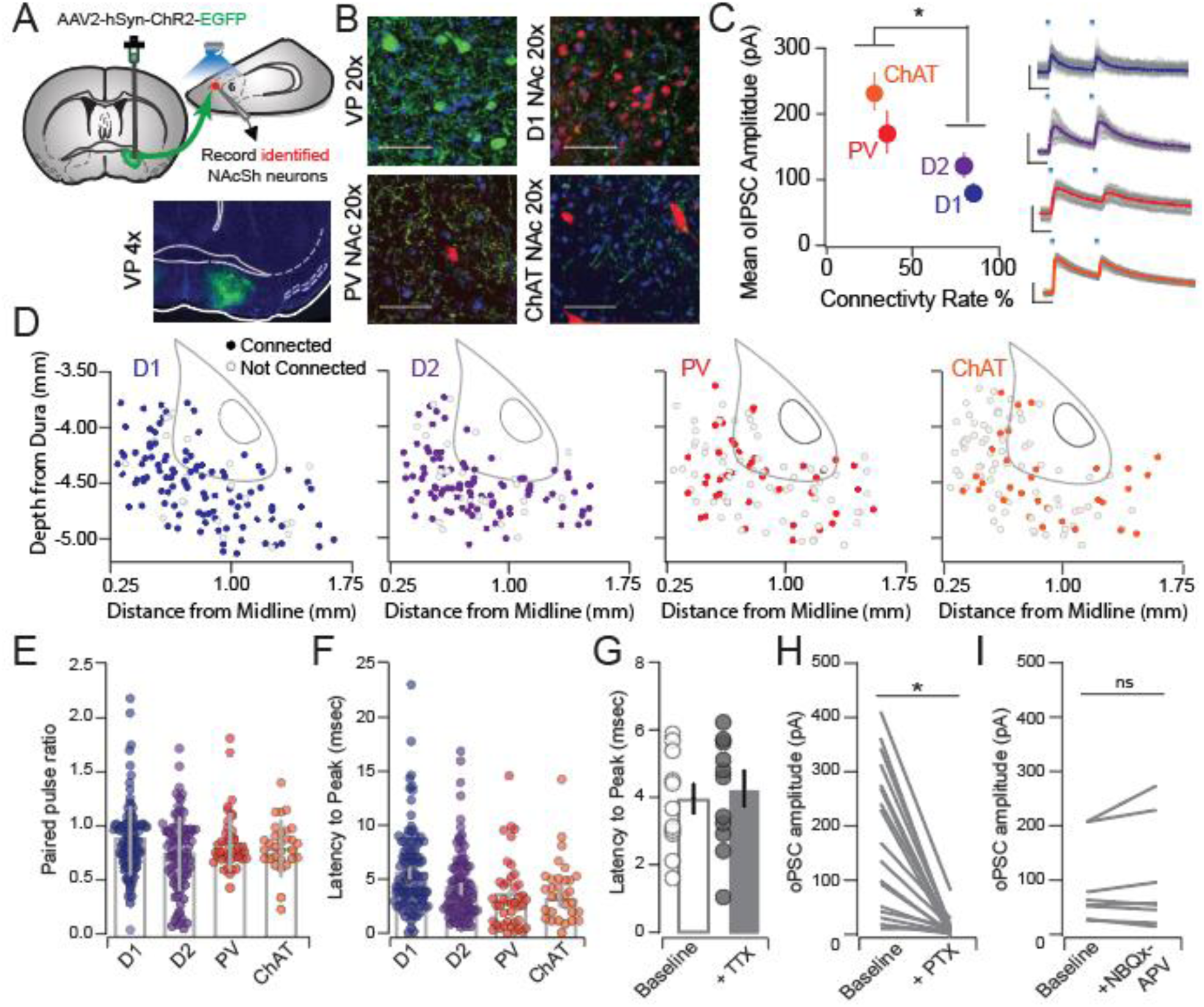
VP neurons make inhibitory, monosynaptic contacts onto NAcSh neurons. **(A)** ChR2 was injected to VP in reporter mice. **(B)** 20x confocal image of ChR2-infected cells in the VP, ChR2-labelled terminals in the NAcSh in D1-To mice, PV-To mice and ChAT-Cre mice injected with DIO-mCherry in the NAcSh. Scale bar = 50 µM. **(C)** Quantification of oIPSCs in D1-MSNs (n=118 cells / 11 mice; 84%, 84.74 ± 8.11 pA), D2-MSNs (n=117 cells / 12 mice; 79%, 114.28 ± 12.23 pA), PVs (n=112 cells / 12 mice; 30%, 163.28 ± 30.70 pA), and CINs (n=104 cells / 10 mice; 28%, 228.65 ± 27.25 pA, Connectivity rate: Pearson x^2^ ratio = 106.24, p<0.001). Right = representative IPSC traces from each cell type, scale bar = 20 msec, 50 pA. **(D)** Distribution of recorded neurons throughout the NAcSh for each neural subtype. **(E)** There was no difference in PPR (D1-MSNs = 84.74 ± 8.11, D2-MSNs = 114.28 ± 12.23, PVs = 163.28 ± 30.70, and CINs = 228.65 ± 27.25; F_3_=2.81, p = 0.04, no significant differences by Bonferroni post-hoc test), or **(F)** latency from light onset to oIPSC peak across geneotypes (D1-MSNs: 4.18±0.42, D2-MSNs: 3.77±0.36, PVs: 3.37±0.52, CINs = 3.26±0.34; F_3_=1.81 p=0.14, no significant differences by Bonferroni post-hoc test). **(G)** Latency to peak was not affected by TTX (Baseline = 16.46 ± 0.69 msec, n=11, TTX = 16.15 ± 0.74 msec, t_11_=0.3113, p = 0.759). **(H)** Currents were abolished by PTX (aCSF = 206.92 ± 37.61 pA, PTX = 24.82 ± 9.88 pA, t_11_=5.415, p=0.0002) **(I)** but not by co-application of NBQX and APV (aCSF = 324.25 ± 250.89 pA, PTX = 357.86 ± 277.72 pA, t_7_=1.215, p=0.264). *p<0.001.

While, these experiments establish that the VP makes broad inhibitory, mono-synaptic connections onto NAcSh neurons, these post-synaptic neurons make extensive local connections. Therefore, poly-synaptic network effects of stimulation of vArky neurons on NAcSh activity *in vivo* would not be evident in the slice preparation. To address this limitation, we transfected the VP with ChR2 as in our slice experiments, and implanted an optic fiber coupled to a recording array in the NAcSh (Fig 4A,B S5A). We recorded 32 multi units and 23 single units from 6 freely moving mice. Single units were classified as putative MSNs (pMSNs) or interneurons (pINs) according to firing rate, waveform peak-valley duration and coefficient of variation (CV) in firing rate. Among the recorded single units, 9 putative MSNs and 14 putative interneurons were identified (Fig 4C, S5B-D, MSN FR = 0.40 ± 0.11 Hz, P-V = 462.8 ± 27.96 µs, CV = 1.77 ± 0.254, IN FR = 3.53 ± 1.20 Hz, P-V = 242.01 ± 26.27 µs, CV = 1.23 ± 0.17). Overall, 53.62% of units were light-modulated, with the vast majority of these units (91.89%) exhibiting a significant *decrease* in firing rate in response to VP terminal activation; a modest increase in firing rate was observed in 3 multi-units in response to a 1 sec or 15 msec light pulse (Fig 4D-H, S5D-H). There was no difference in the proportion of light-modulated units between multi-units, pMSNs or pINs, nor was there a significant difference in the magnitude of inhibition between these unit classes (Fig 4H). These results demonstrate that activation of VP terminals in the NAcSh is sufficient to robustly inhibit NAcSh firing in vivo.

**Figure 4.**
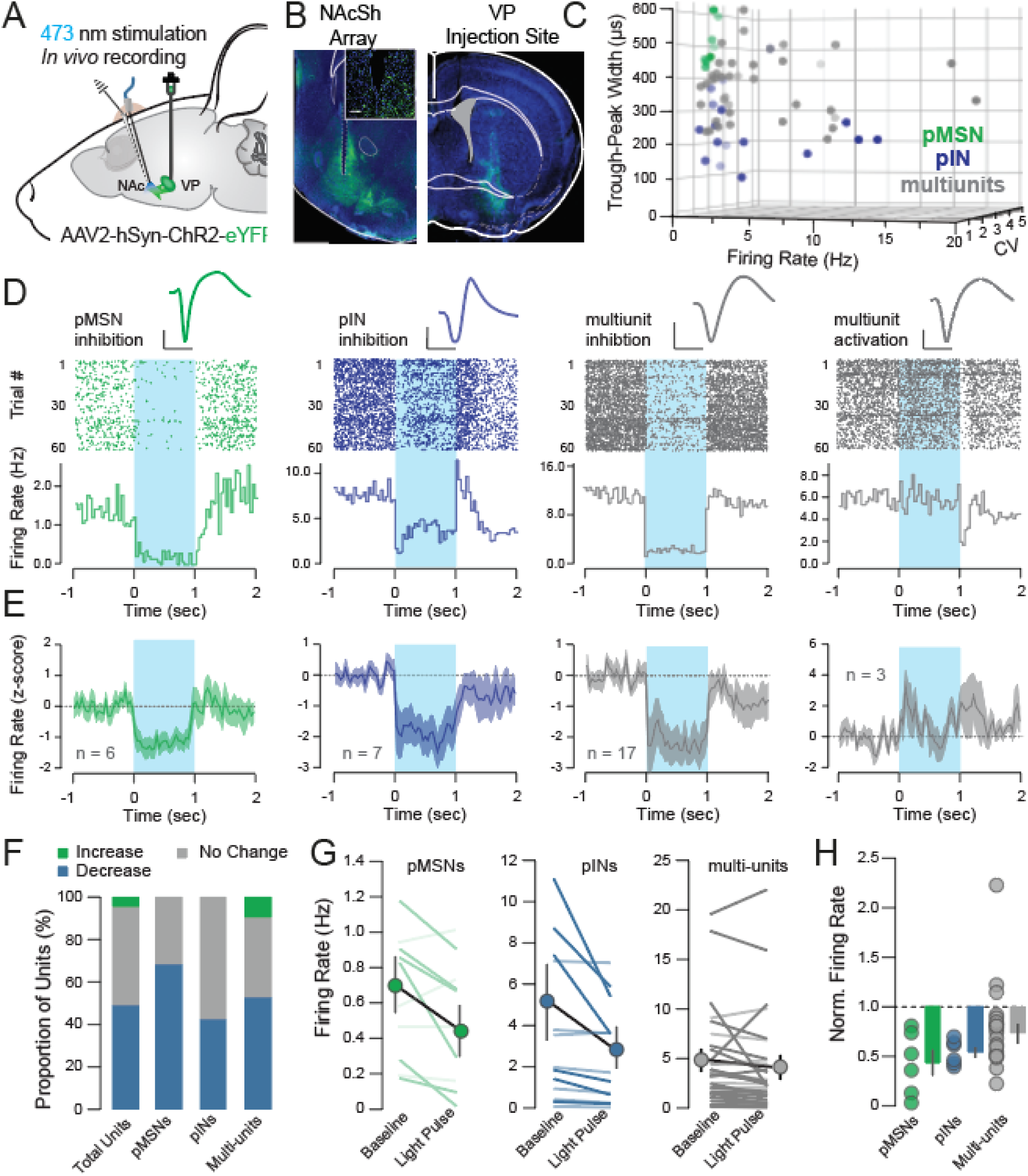
Activation of vArky terminals inhibits NAcSh firing in vivo. **(A)** ChR2 was injected in the VP, a recording array with optic fiber was implanted in the NAcSh. **(B)** 4x images of NAc and VP; inset: 20x confocal image of array site. **(C)** Units were classified into clusters by peak-to-valley width, firing rate and CV (n = 9 pMSNs, 14 pINS, 32 multi-units from 6 mice, see Fig S5). **(D-E)** Representative waveform, raster plot and PSTH of single neuron examples and averages of light-modulated units over 60 trials; 1 sec light pulse is indicated. Scale bar = 500µs, 50µV. **(F)** Proportion of light-modulated units by neural subtype. **(G)** Firing rate immediately preceding and after light pulse are shown for each cell-type; dark lines: light-modulated units, light lines: non-significantly modulated units. **(J)** Normalized firing rate of all light-modulated units, relative to baseline is shown (multiunits = 0.66 ± 0.05, pMSNs = 0.46 ± 0.13, pINs = 0.59 ± 0.05, F_2_ = 1.789, p =0.185).

### Endogenous vArky calcium is associated with duration of reward consumption

Given our findings that vArky neurons inhibit NAcSh neurons in vivo and that NAcSh inhibition drives reward consumption (Fig 1; O’Connor et al., 2015; Roitman et al., 2010), we hypothesized that vArky neurons would be active during reward consumption, and the extent of this activity at reward consumption onset would predict length of ensuing lick bouts. To test this, we used a retrograde viral strategy to express gCaMP6s exclusively in vArky neurons and recorded bulk calcium activity using fiber photometry (Fig 5A-C). We aligned calcium activity to the onset of lick bouts during a free-access task, and determined that, as predicted, fluorescence signal peaked following reward consumption onset; there was no lick-related fluorescence change in GFP-controls (Fig 5D, S6A,C). To determine whether endogenous vArky activity predicted the duration of subsequent reward consumption, we calculated the area under the curve (AUC) of each trial, and compared the upper (GCaMP high trials) and lower quartiles (GCaMP low trials) for each subject (Fig 5E, S6A). Consistent with our hypothesis, duration of lick bouts was significantly *longer* during high GCaMP trials relative to low GCaMP trials (Fig 5F). This relationship was observed across every subject, and was not observed in GFP controls (Fig S6B-E). To further evaluate the significance of this behavioral difference between high- and low-GCaMP trials, we performed a permutation analysis in which we shuffled the assignment of high- and low-trials 2000 times, and detected an average difference of 0.6%, while the observed difference in lick behavior between high- and low-GCaMP trials was 42.6% (Fig 5G). Our hypothesis predicts that excitatory responses in vArky neurons would precede NAcSh inhibition. We recorded NAcSh activity with the fiber photometry modality, and confirmed that a decrease in NAcSh activity did not occur prior to reward consumption on-set (Fig S7A-C).

**Figure 5.**
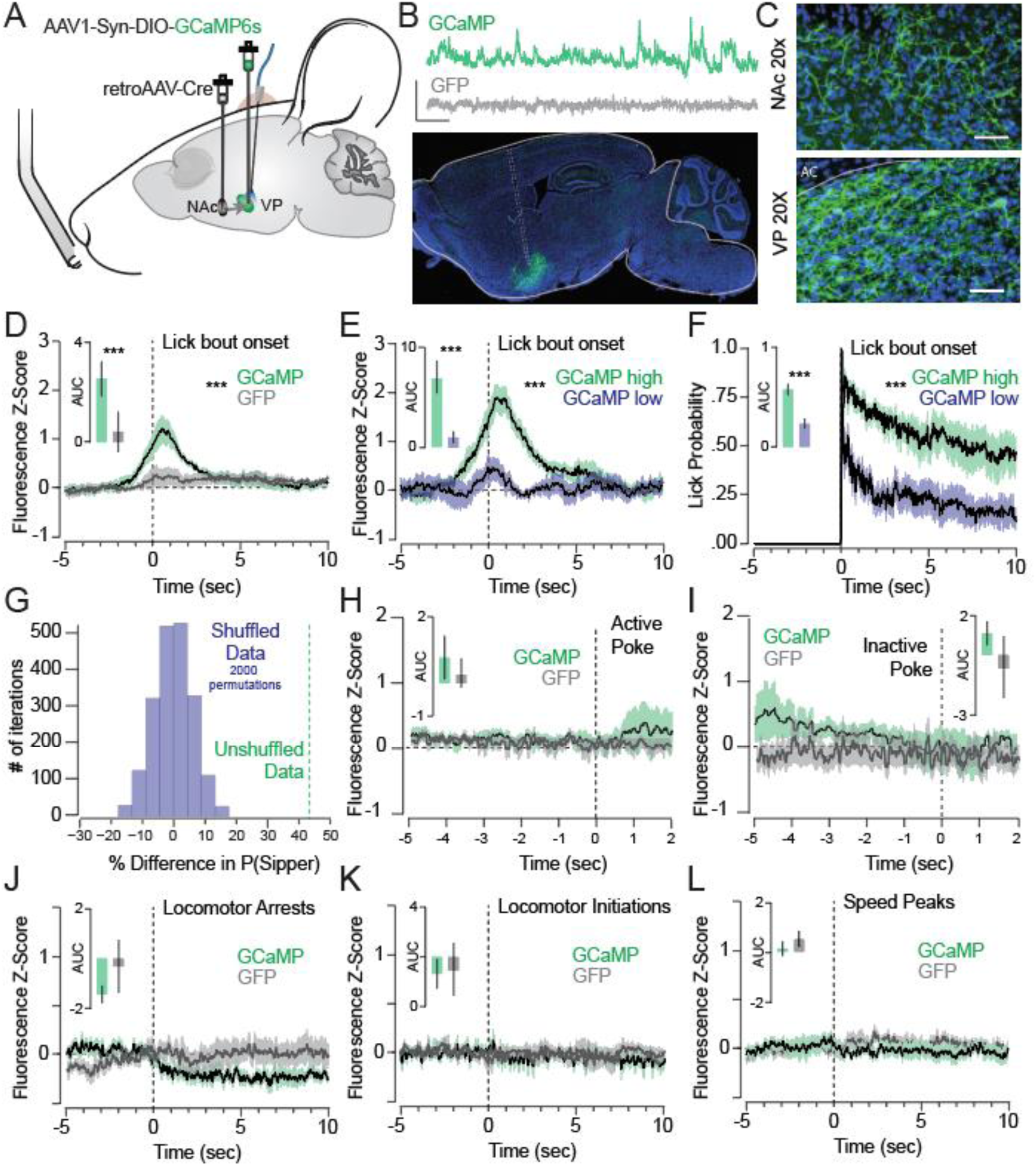
High endogenous vArky calcium activity at reward consumption onset is associated with longer duration of reward consumption. **(A)** retroAAV-Cre was injected to the NAcSh, AAV-DIO-GCaMP6s was injected in the VP, an optic fiber was implanted in the VP. **(B)** Representative traces of Ca^2+^ signal acquired from optic fiber implanted in VP (scale bar=20s, 1 z-score). **(C)** 20x confocal images of cell bodies in the VP and terminal fields in the NAc (scale bar=50 µM). **(D)** Combined fluorescence responses from all mice, aligned to onset of lick bout (AUC GCaMP: 2.43±0.75, AUC GFP: 0.52±0.71, F=7.213, p=0.016, n=9 mice/group). **(E)** Fluorescence responses in GCaMP mice, split into upper and lower quartiles based on AUC (44 trials/condition, AUC high trials: 5.76±0.575, AUC low trials: 2.37±0.4911, RM ANOVA, F=13.86, p < 0.0001). **(F)** Licking behavior aligned to bout onset for high- and low-GCaMP signal trials (AUC_high signal trials_: 5.76±0.57, AUC_low signal trials_: 2.37±0.49, F_High•Low_=20.08, p<0.001). **(G)** Permutation analysis: distribution of mean difference in lick bout probability between high- and low-VP signal trials with 2000 permutations of shuffled trials (P_shuffled_(LickBout)VP-High trials – P_shuffled_(LickBout)VP-Low trials = 0.6%), mean difference in real data shown in green (P_real_(LickBout)VP-High trials – P_real_(LickBout)VP-Low trials = 42.6%). **(H-I)** GGaMP and GFP signal aligned to active (**H**, mean AUC GCaMP: 0.78±0.66, GFP: 0.30±0.46) and inactive nose pokes (**I**, mean AUC GCaMP: 1.09±0.54, GFP: −0.65±1.56) in an operant task (F_virus_ = 0.58, p=0.46, F_active v inactive_=1.83 p=0.20). **(J-L)** Fluorescence responses aligned to locomotor arrests (mean AUC GCaMP: −1.467±0.2994, GFP: −0.3198±1.071, t_16_=1.127, p=0.27), locomotor initiations (n=704/677 events, AUC GCaMP: −0.818±0.638, GFP: - 0.619±1.210, t_16_=0.150, p=0.88) and peaks in locomotor speed (n=1309/1221 events, mean AUC GCaMP: 0.1294±0.2945, GFP: 0.4977±0.3022, t_16_=0.873, p=0.396). ***p<0.001.

Finally, we determined that vArky activity was not related to reward seeking per se by introducing a requirement for a nose poke before reward availability. There was no significant change in GCaMP signal preceding or immediately following the seeking nose-poke, prior to reward consumption, or around inactive nose pokes (Fig 5H,I). We performed three additional analyses to rule out the possibility that the reward consumption responses were driven by motor-related activity in vArky neurons that may occur during reward approach. We first aligned the photometry signal to locomotor arrests. In contrast to the *increase* in signal around reward consumption, there was no significant change in the fluorescence signal with locomotor arrests not associated with reward consumption (Fig 5J). There was also no significant change in GCaMP signal related to locomotor initiations or to peaks in ambulatory velocity (Fig 4K,L). These results confirm that the magnitude of endogenous calcium response in vArky neurons upon reward consumption onset is associated with the duration of ensuing lick bout.

### Activation of vArky neurons promotes reward consumption

Finally, we hypothesized that activation of vArky neurons would promote reward consumption. We applied closed-loop optogenetic stimulation to drive vArky activity at 20 Hz during reward consumption in a free-access paradigm (Fig 6A, S8A,B). Closed-loop stimulation of vArky terminals significantly increased the total drinking time (214%, Fig 6B). This effect was not due to changes in approach behavior, as measured by number of bouts (Fig 6C), but was driven by an increase in mean bout length (Fig 6D); there was also a significant shift in the distribution towards longer drinking bouts with ChR2 stimulation (Fig 6E). However, open-loop optogenetic vArky stimulation applied randomly throughout the session stimulation did not increase drinking time or number of lick bouts, confirming that vArky stim does not drive approach behavior (Fig S8C D), but rather promotes on-going reward consumption.

**Figure 6.**
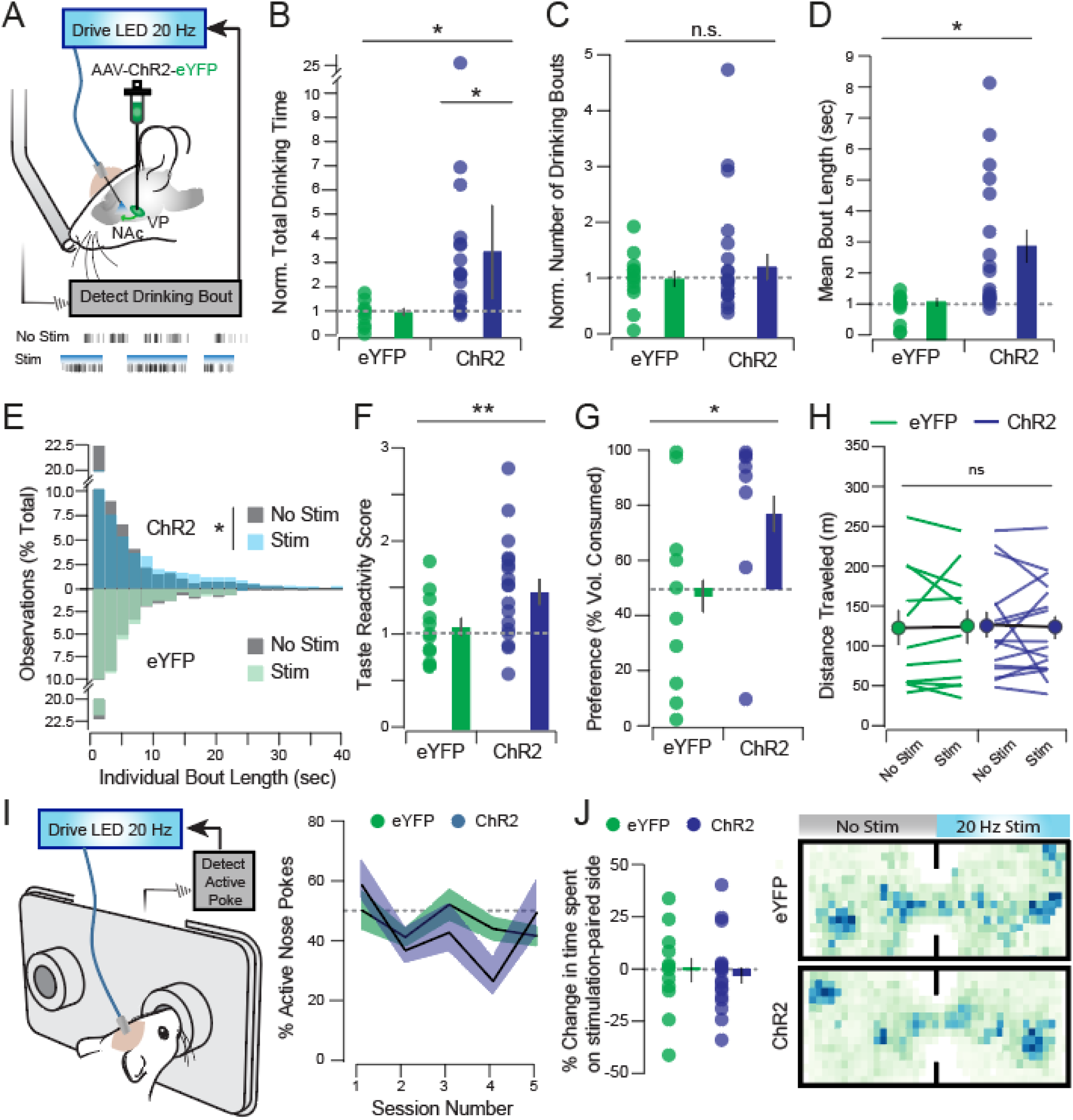
Closed-loop activation of vArky terminals promotes reward consumption. **(A)** Closed-loop optogenetic activation of VP terminals in NAcSh during reward consumption **(B)** Stimulation-induced change from baseline in total drinking time (eYFP: 0.95±0.14, ChR2: 3.84±1.32, F_virus•stim_=5.44, p=0.027) **(C)** number of drinking bouts (eYFP: 0.98±0.14, ChR2: 1.37±0.25, F_virus•stim_=0.42, p=0.840) and **(D)** mean bout length (eYFP: 0.91±0.13, ChR2: 2.83±0.52, F_virus•stim_=6.49, p=0.017). **(E)** Histogram distribution of lengths of each individual bout across all subjects (eYFP_NoStim/Stim_ n=564/558, ChR2_NoStim/Stim_ n=1331/1001). **(F)** Stimulation-induced change in orofacial taste reactivity (eYFP_NoStim_: 3.88±0.46, eYFP_Stim_: 3.99±0.27, p=0.781, ChR2_NoStim_ = 3.36±0.35, ChR2Stim: 4.77±2.05, F_virus•stim_=7.69, p=0.007). (G) Preference for stimulated bottle in a two-bottle choice task (eYFP: 46±11%, ChR2: 74±12%, x^2^=5.05, p=0.025). **(H)** Distance traveled in an open field task (eYFP_NoStim_: 121.62±22.63, eYFP_Stim_ : 122.15±21.75, p=0.96, ChR2_NoStim_: 124.48±16.22, ChR2_Stim_: 121.83±14.18, F_stim•virus_=0.050, p=0.825). **(I)** Proportion of active to inactive nose pokes over 5 days of operant training (eYFP: 44.16±2.08%, ChR2: 41.17±3.47%). **(J)** Preference for the stimulation-paired side of a chamber in a real-time place preference task (eYFP: −4.36±4.32%, ChR2: −1.13±5.83) and representative heatmaps. *p<0.05, **p<0.01. All data mean±SEM.

Reward consumption is driven by dissociable processing of “wanting” (motivation for reward) and “liking” (hedonic value of reward; Berridge, 2000). The selective increase in bout length without a change in the approach behavior indicates stimulation of VP terminals in the NAcSh increased the palatability or hedonic value of reward, rather than changed motivation for reward per se (Boughter et al., 2007; Strickland et al., 2018). We tested this explicitly in two additional tasks. We first measured orofacial reactivity in response to a 10% sucrose solution. vArky stimulation increased hedonic tongue protrusions during stimulation relative to un-stimulated trials (Fig 6F), arguing in favor of increased hedonic value of reward. Further, when two identical liquid rewards were presented in a two-bottle choice paradigm, significantly more mice exhibited a preference for the bottle paired with optogenetic vArky stimulation relative to EYFP controls (Fig 6G).

Optogenetic vArky stimulation did not affect locomotor activity in either eYFP- or ChR2-injected mice (Fig 6H), arguing that the increase in reward consumption is not due to inhibition of locomotor activity. Non-selective optogenetic or electrical stimulation of the VP is reinforcing (Panagis & Spyraki, 1996; Tooley et al., 2018), raising the possibility that the increase in reward consumption arose because vArky stimulation alone is rewarding. To test this possibility, we performed an operant task, and found that neither eYFP- or ChR2-injected mice showed a preference for the active port which delivered vArky stimulation, over an inactive port that had no consequence (Fig 6I). Moreover, in a real-time-place-preference task, eYFP- and ChR2-injected mice exhibited no preference for the stimulation-paired side of a chamber relative to the unpaired side (Fig 6J). These results establish that vArky stimulation in the absence of reward is not reinforcing, and that activation of the vArky pathway promotes reward consumption by amplifying the hedonic value of reward.

## DISCUSSION

While classic models of information flow in the basal ganglia posit that the VP is exclusively and output of the NAc, we show that a subpopulation of VP neurons provides dense, inhibitory inputs throughout the NAcSh. We further show that activity of this vArky pathway promotes reward consumption by enhancing reward value, providing a novel circuit mechanism by which the VP orchestrates mesolimbic activity to mediate reward processing.

### Functional organization of the vArky pathway

Classical models of the basal ganglia posit that the NAc integrates inputs from cortical and limbic areas and projects downstream to the ventral pallidum, the so-called “limbic motor interface”, that transmits information to downstream motor structures for expression of behavior (Heimer et al., 1982). However, early tracing studies have suggested that thin pallidofugal axons traversing the NAc originate in the VP (Haber et al., 1985). Moreover, an analogous arkypallidal projection has been described in the dorsal basal ganglia, which originates in the GPe and provides a rapid source of inhibition to the striatum to allow flexible action selection (Mallet et al., 2012, 2016). We confirm that a subset of VP neurons sends dense, web-like processes throughout the NAcSh, and that these vArky neurons form inhibitory synapses onto both NAcSh MSNs and interneurons. vArky neurons project to both MSNs and INs, although we observe relatively higher connectivity rates onto MSNs (Fig 2,3,S2). This anatomical organization is analogous to the projection that has been described in the dorsal basal ganglia (Glajch et al., 2016; Hernández et al., 2015), and argue for common core organizational principles between the dorsal and ventral Arky neurons.

### Modulation of the NAcSh by the vArky pathway

The NAcSh is a crucial structure for affective processing and generating motivated responses to rewards (Floresco, 2015). Both D1- and D2-MSNs in the NAcSh send inhibitory projections to VP, lateral hypothalamus and VTA, and inhibition of the NAcSh is thought to release these downstream structures from tonic inhibition (Creed et al., 2016; O’Connor et al., 2015; Smith et al., 2011; Stratford, 2005). Consistent with this hypothesis, NAcSh inhibition is both necessary and permissive for reward consumption. NAcSh units robustly decrease firing during consumption of palatable rewards (Fig 1A-D; Cheer et al., 2005; Janak et al., 2004; Krause et al., 2010; Nicola et al., 2004a, 2004b; Roitman et al., 2010; Taha & Fields, 2006). The high proportion of inhibited NAcSh units argues against the idea that one NAcSh cell type is specifically implicated; and studies that have measured both populations with photometry or optotagging reveal consumption-related inhibitory responses in both D1- and D2-MSNs, although these responses are more prevalent in D1-MSNs (London et al., 2018; O’Connor et al., 2015). NAcSh inhibition is *necessary* for reward consumption, as infusion of glutamate agonists to the NAcSh reduce reward consumption, and optogenetic activation of accumbal units can abort on-going reward consumption (Krause et al., 2010; O’Connor et al., 2015). This inhibition is *sufficient* to promote reward consumption, since pharmacological or optogenetic inhibition of the NAcSh robustly increases reward consumption in the absence of metabolic need (Fig 1E-H, O’Connor et al., 2015; Stratford, 2005; Zhang et al., 2003). Despite the importance of this NAcSh inhibition for reward consumption, the substrate of this pause in NAcSh firing is not completely understood.

Decreased excitatory drive is one potential substrate of this NAcSh inhibition. Activity of excitatory projections from thalamus, hippocampus and amygdala to NAcSh is reduced during reward consumption, while pharmacological or optogenetic inhibition of these excitatory inputs produces appetitive motivation (Faure et al., 2008, 2010; Reed et al., 2018; Reynolds & Berridge, 2003). However, inhibition of NAcSh glutamate does not amplify hedonic responses to reward (Faure et al., 2010; Reynolds & Berridge, 2002; Richard et al., 2013). Instead, activation of GABAergic receptors within the NAcSh induces this effect. While local collaterals between GABAergic NAcSh neurons is one source of inhibition, our results establish that the VP is an important exogenous source of GABAergic input that regulates NAcSh firing and reward-value. The lateral inhibition is interesting in light of our in vivo results; a minority (3 muli-units) of NAcSh neurons exhibited increased firing in response to vArky terminal activation. This is likely due to release of lateral inhibition from inhibited MSNs and INs, given the extensive lateral connections within the NAcSh. Future studies will establish whether these activated units represent a distinct functional subclass or play a unique role in reward processing, since subsets of NAcSh units show increased firing in response to reward (Fig 1B, Ambroggi et al., 2011).

### Activity of vArky neurons predicts and drives reward consumption

The VP is unique in that firing activity tracks hedonic value of reward which is dissociable from reward-related motivation; neuronal firing rates increase as a function of experienced and expected reward value, and decrease with devaluation (Ottenheimer et al., 2018; Tachibana & Hikosaka, 2012; Tindell et al., 2006, 2009). Moreover, consumption-related neural responses are more prevalent in the VP relative to the NAc (Ottenheimer et al., 2018) and VP neural responses to consumption precede consumption signals in the NAc (Ottenheimer et al., 2018; Richard et al., 2016). The vArky pathway is a compelling mechanism by which early reward-related responses in VP can directly promote the inhibition in NAcSh firing which is necessary for reward consumption.

Consistent with this hypothesis, calcium activity in these vArky neurons occurs prior to reward consumption, showing anticipatory ramping during un-cued reward approach. This signal does not reflect reward seeking, per se, since there was no calcium activity in response to active instrumental responses during an operant task. Instead, our results reveal that the magnitude of peak in calcium activity at reward consumption onset was associated with the length of the ensuing lick bout; confirming that early activity of vArky neurons could predict subsequent reward consumption. While the VP also encodes motor activity independent of reward; calcium activity did not reflect locomotor arrests or ambulatory activity outside of reward consumption. These results confirm that vArky activity calcium activity is selective to reward consumption and suggest a causal role for vArky neurons in sustaining reward consumption.

We establish this causal role of vArky neurons in sustaining reward consumption in a value-dependent manner. vArky activation altered the microstructure of reward consumption by selectively increasing the length of individual bouts (Fig 6D,E) which is one index of hedonic value (Boughter et al., 2007; Strickland et al., 2018). Moreover, vArky stimulation increased hedonic orofacial reactions (Fig 6F), which reflects affective responses to reward independent of motivation (Berridge, 2000). Finally, vArky stimulation drove preference for a stimulation-paired reward given two otherwise identical rewards in a two bottle choice task (Fig 6G), further confirming a role for vArky neurons in modulating reward value. Interestingly, while decreased excitatory drive to the NAc has been shown to drive reward approach and consumption (Reed et al., 2018), open-loop vArky stimulation did not increase reward approach, suggesting that vArky does not initiate, but rather maintains this inhibition in a value-dependent manner. Moreover, open-loop stimulation did not alter bout length or drinking time. This indicates that vArky activity modulates reward consumption on short time scales, rather than inducing long-term effects on inhibitory tone in the NAcSh. Together, these results establish a selective role for the vArky pathway in promoting reward consumption as a function of reward value.

### Concluding remarks

Our results establish that a unique subpopulation of VP neurons, the vArky neurons, project to and inhibit the NAcSh, and that this pathway sustains reward consumption by enhancing hedonic value of reward. Integration of this pathway into models of the basal ganglia is necessary to understand how value and motivation for reward is encoded within these nuclei, and how coordinated activity across this network orchestrates reward-seeking behavior.

## Supporting information

Supplemental Video cleared hemisphere

## ACKNOWLEDGEMNTS

We thank AV Kravitz for expertise and assistance with in vivo electrophysiology and fiber photometry experiments, and MC Stander for excellent technical help. Research was supported by internal funds from Washington University in St. Louis, the Brain and Behavior Research Foundation (NARSAD Young Investigator Grant #27197 to MCC.), National Institutes of Health National Institute on Drug Abuse (#R21-DA047127, R01-DA049924 to MCC), Whitehall Foundation Grant (#2017-12-54 to MCC.), and Rita Allen Scholar Award in Pain (to MCC.).

## AUTHOR CONTRIBUTIONS

YMV, KA and MCC wrote the paper with input from all authors. Anatomical experiments were performed by: YMV, JRT, HS and TE; YMV and MCC performed patch clamp physiology, JRT and OU performed in vivo electrophysiology, JRT and EG performed fiber photometry, YMV, JRT, EC and KA performed behavioral experiments. Data was analyzed by YMV, JRT and MCC. Work was supervised by CL and MCC.

## EXPERIMENTAL PROCEDURES

### Subjects

Adult male and female mice on a C57BL/6J background were bred in-house from stock lines obtained from Jackson Labs (D1-Cre, 129-Tg(Drd1-cre)120Mxu/Mmjax; PV-Cre, 129P2-Pvalb<tm1(cre)Arbr>/J; ChAT-Cre, 129S6-Chattm2(cre)Lowl/J; D1-To, B6.Cg-Tg(Drd1a-tdTomato)6Calak/J; PV-To, 6-Tg(Pvalb-tdTomato)15Gfng/J JAX# 027395, or wildtype C57Bl6/J) or MMRC repository (A2a-Cre; Tg(Adora2a-cre)KG139Gsat). Mice were group housed (2 – 5/cage, single sex) in a vivarium that was controlled for humidity, temperature, and photoperiod (12 : 12; lights on at 7 am and off at 7 pm). Mice had ad libitum access to food and water throughout the experiment. All procedures were approved by the Institutional Animal Care and Use Committee at Washington University in St. Louis and the University of Maryland and were conducted in accordance with the *Guide for the Care and Use of Laboratory Animals*, as adopted by the NIH.

### Stereotaxic Surgery

Anesthesia was induced and maintained with isoflurane at 5% and 1.5%, respectively. Mice were placed in a stereotaxic frame and craniotomies were performed. For optogenetic, electrophysiology, photometry and behavioral experiments, purified and concentrated adeno-associated viruses were injected into the NAc (coordinates relative to Bregma : AP: +1.7mm, ML: −0.6mm, DV: −4.5mm), the VP (coordinates relative to Bregma, AP: +0.2 mm, ML: ±1.5 mm, DV: 4.6 mm). GCaMP6s virus (pAAV.Syn.DIO.GCaMP6s.WPRE.SV40) was obtained from Addgene, Rabies-EGFP virus (EnvA G-deleted Rabies-EGFP) was obtained from the Salk Vector Core, all other viruses (rAAV2Retro-Cre, AAV5-EF1a-DIO-hChR2(H134R)-EYFP, AAV-hSynhChR2(H134R)-EYFP, AAV-CAG-FLEX-GFP, AAV-CA-FLEX-RG, AAV-EF1a-FLEX-TVAmCherry, AAVhSyn-eArch3.0-EYFP, AAV-EF1a-DIOmCherry) were obtained from the University of North Carolina Vector Core Facility. Injections were carried out using graduated pipettes broken back to a tip diameter of 10-15 μm, at a rate of 0.05 μl / min for a total volume of 0.25-0.5 μl; pipettes were left in situ for an additional 8 minutes to allow for the diffusion of virus; 28 days was the minimum viral expression time. For behavioral experiments, optic fibers were implanted bilaterally over the NAcSh or VP and secured with 3 skull screws and dental cement. Animals recovered for at least 1 week before behavioral training, and only mice with verified infection sites and fiber placements were included in the analyses.

### Neuroanatomical Tracing and Histology

#### Retrograde ChR2 tracing

250 nL of retroAAV-Cre was injected to the NAcSh, and 300 nL AAV2-DIO-hSyn-ChR2-eYFP was injected in the VP. Following 28 days expression, tissue was harvested, and ChR2 signal was amplified by immunostaining. Briefly, free floating brain sections were permeabilized in 3 washes of PBST (PBS with 3% TritonX) for 10 min each at room temperature, blocked for 1 hour in 5 % normal donkey serum in 3% PBST and then incubated in chicken anti-GFP (Invitrogen, A10262, 1:1000) overnight at 4°C. Brains were washed in 3 times in 3% PBST (10 min) and incubated in goat anti-chicken Alexa Fluor 488 (Invitrogen, A11039, 1:1000) in blocking buffer at room temperature for 2hrs. After 3 additional 10 min PBST washes and 3 washes in PBS, sections were treated with SudanBlack fluorescence quencher (Sigma) and were coverslipped and mounted using VectaShield mounting solution containing DAPI (Vector Laboratories). Intensity of fluorescence signal was quantified across 3 z-stacks from separate serial sections (20x, 500µM^2^) in each of the following structures in 6 mice (2M, 4F): nucleus accumbens shell (NAcSh), ventral pallidum (VP), medial prefrontal cortex (mPFC), orbitofrontal cortex (OFC), basolateral amygdala (BLA) lateral habenula (LHb), mediodorsal thalamus (MDThal), subthalamic nucleus (STN), ventral tegmental area (VTA).

#### Immunolabeling and clearing of hemibrains by iDisco and light sheet imaging

The iDISCO+ protocol was followed as described in (Renier et al., 2014, 2016). Hemibrains were immunolabeled with Chicken anti-GFP (ThermoFisher, Cat# A10262) at 1:200 during a 5 day incubation at 37°C. Secondary antibody AlexaFluor 647-conjugated goat anti-chicken (ThermoFisher #A21449) was used at 1:200 iDisco-processed hemibrains were imaged by light sheet microscopy (Miltenyi Biotec Ultramicroscope II) using their 4x objective. All images were acquired using The Right or Left light sheets (3 sheets per side) with dynamic focusing (10 positions) enabled. For the Dynamic Focus Processing the Blend algorithm was used.

#### Retrobeads

300 nL of Red Retrobeads and Green Retrobeads (both Lumafluor) were injected into the NAcSh (coordinates from bregma: 1.7AP, 0.6ML, −4.5DVmm) and VTA (coordinates from bregma: −3.2AP, 0.8ML, −4.3DVmm), respectively. Tissue was harvested 10 days following injection, and retrobead-expressing cells were quantified through 8 serial sections through the VP from 8 mice (5F, 3M).

#### Fluorescent In Situ Hybridization

Mice were injected with green retrobeads in the NAcSh (see above) 10 days after injection, mice were euthanized and fresh tissue was frozen on dry ice, stored at −80°C and sectioned with a cryostat (Leica, CM3050) at 20 µM. Tissue was processed using RNAscope® Technology in situ hybridization assay according to provided protocols (ACDBio). Briefly, slides were transferred from −80°C and fixed with formalin, dehydrated with increasing concentrations of ethanol and pretreated with provided protease solution before hybridization with probes targeted against Slc32A1 and Slc17A6 mRNA. Following 2 h of hybridization at 40°C, sections were washed and fluorescence was amplified with provided buffer solutions. Sections were incubated with DAPI before cover-slipping for visualization.

#### Pseudotyped rabies tracing

Rabies “helper virus” (AAV5-CAG-DIO-RG, AAV5-EF1a-DIO-TVA-mCherry, 250 nL in a 1:1 mixture, University of North Carolina) was injected to the NAcSh in the following Cre-driver mouse lines: ChAT-Cre, PV-Cre, D1-Cre, A2a-Cre. After 14 days expression, EnvA G-Deleted Rabies-eGFP (Salk Vector Core) was injected to the same coordinates. Following 7 days expression, tissue was harvested, and serial sections were taken throughout the NAc and VP. Starter cells were quantified in the NAcSh, defined as cells-co-expressing mCherry and EGFP; EGFP-positive cells were counted throughout the rostral-caudal extent of the VP using threshold detection function in ImageJ (Schindelin et al., 2012).

#### Perfusion

Mice were deeply anesthetized with 5% isofluorane and intracardially perfused with 1X phosphate buffered saline (PBS) and then with 4% paraformaldehyde in PBS 1X. Brains were frozen in cooled isopentane (−50°C) and stored at −80°C. Serial coronal or sagittal sections were cut at 40 μm with a cryostat (Leica CM3050) collected in PBS and stored at 4°C.

#### Imaging

Slides were initially scanned with an upright fluorescent slide-scanning microscope (Leica DM6 B) at 5X and visualized with Leica Application Suite X software to identify areas of interest. For fluorescent immunohistochemistry experiments, sections were imaged with a confocal laser-scanning microscope (Leica, DMI4000 B) with a 20X air objective. The brightness and contrast of images were adjusted to optimize visualization, and adjustments were applied uniformly across each image. For rabies tracing experiments, 20x images acquired from the Leica slide-scanning microscope; regions of interest were manually delineated based on the atlas of Paxinos and Watson (Franklin, Keith, 2019), and subsequently imported to ImageJ for cell counting.

### Patch Clamp Electrophysiology

Whole-cell patch-clamp recordings were made from coronal slices (220 µm) of the NAcSh. Slices were prepared using a vibratome (Leica VT 2100) in ice cold cutting solution (in mM: 0,5 CaCl2, 110 C5H14CINO, 25 C6H12O6, 25 NaHCO3, 7 MgCl2, 11.6 C6H8O6, 3.1 C3H3NaO3, 2.5 KCl, and 1,25 NaH2PO4) and continuously bubbled with 95% O_2_ and 5% CO_2_. Slices were incubated at 32°C for 30 min in artificial cerebrospinal fluid (aCSF; in mM: 119 NaCl, 2.5 KCl, 1.3 MgCl_2_, 2.5 CaCl_2_, 1.0 Na_2_HPO_4_, 26.2 NaHCO_3_, and 11 glucose) followed by storage at room temperature until electrophysiological recordings were performed. Slices were hemisected and superfused with aCSF at 30 ± 2°C. Neurons were recorded using borosilicate glass pipettes (5-7 MΩ resistance) pulled on a micropipette puller (Narishige PC-100) filled with cesium-based internal solution (in mM: 110 CsMeS, 15 KMeS, 0.02 EGTA, 10 Hepes, 0.5 MgCl_2_, 1.5 MgSO_4_, 4 ATP-Na_2_, 0.3 GTP-Na_3_, 10 C_6_H_12_O_6_, and 10 Phosphocreatine-Na_2_ with 7.3 pH and 289 Osm). Currents were amplified, filtered at 2 kHz, and digitized at 10 kHz using a MultiClamp 700B amplifier and Digidata 1550 (Molecular Devices). Clampex 11.4 software (Molecular Devices) was used for data acquisition. Series resistance was monitored by a hyperpolarizing step of −15 mV for 5 msec every 10 sec; the cell was discarded if the series resistance changed by more than 15%. Neurons were visualized using an Olympus 560X upright microscope; wholefield LED illumination (CoolLED) was used to visualize TdTomato or mCherry (560 nm), and stimulate ChR2-expressing terminals (473 nm, paired 4ms light pulses, 50 ms ISI, 13.7-18.2 mW). All agents were purchased from Sigma Biosciences. Coordinates of recorded cells in the NAcSh were captured at 4x on the patch microscope and registered to the atlas of Paxinos and Watson.

### In vivo electrophysiology

Wildtype mice (3M, 3F) were injected with ChR2 in the VP, and 16-channel nickel-coated recording arrays (MWA-250×250-NiCr-50-16-5 -MOD16-1-G2, 50 µm, MicroProbes for Life Science) coupled to an optic fiber (ThorLabs, constructed in house) were implanted in the NAcSh. Neurophysiological signals were transmitted to an Omniplex neurophysiology system (Plexon Inc) via a multiplexing digital headstage, while 473 nm light was delivered through the optic fiber via a Plexon LED and driver system. Spike channels were acquired at 40 kHz with 16-bit resolution, and band-pass filtered at 150 Hz to 3 kHz before spike sorting. Local field potential signals were simultaneously digitized at 1Khz. Single and multiple units were discriminated using principal component analysis (Offline Sorter; Plexon), using MANCOVA analyses to determine if single unit clusters were statistically distinct from multi-unit clusters. Single units exhibited a mean 2D J3 = 3.41, Daves-Boulin = 0.272, and p value by MANCOVA = 0.00801. Single units were classified into putative MSNs (pMSNs) and putative interneurons (pINs) based on peak-valley duration: pMSNs 462.8 ± 27.96, pINs 242.01 ± 26.27,multi-units 360.1 ± 19.49, CV: pMSNs 1.77 ± 0.25, pINs 1.23 ± 0.17, multi-units 1.49 ± 0.14, Firing Rate: pMSNs 0.40 ± 0.11, pINs 3.53 ± 1.20, multi-units 4.12 ± 0.91. Mice were recorded three times, and the recording with the largest number of channels with unit activity was chosen for analysis.

To identify light-modulated units, we exported the average peri-event-histogram (PSTH) of spiking of each unit around light onset in 10 msec bins. To test whether a neural response of an isolated NAcSh unit was significantly modulated by activation of local VP terminals, paired t-tests were performed on the average firing activity of each isolated unit in the 50 msec period during light onset relative to activity during the 1 sec baseline period immediately preceding light onset. A unit was determined to be light-modulated if the difference between the average activity between baseline and light-onset was statistically significant. We used post-morten histology and modulation of the LFP by blue light to confirm our recording sites; if there was no significant modulation of LFP by blue light stimulation, units recorded on the same channel were excluded from further analysis (resulted in exclusion of 3 multi - units from 2 mice). Autocorrelelagrams were performed to exclude duplicate units.

### Fiber Photometry

Wildtype mice (9M, 11F) were injected with AAVDJ-hsyn-GCaMP6s-eYFP (NAcSh measurements) or retroAAV-Cre into the NAcSh (vArky measurements, 250 µL, coordinates from bregma: AP: +1.7mm, ML: −0.6mm, DV: −4.5mm). For vArky measurements, AAVDJ-hsyn-DIO-GCaMP6s-eYFP was injected to the VP (300 µL, coordinates from bregma: AP: +0.6mm, ML: −1.5mm, DV: −4.65mm (Franklin, Keith, 2019); an additional 9 mice (5M, 4F) were injected with DIO-GFP as controls. An optic fiber (200 µM core, 0.39 numerical aperture, ThorLabs) was implanted unilaterally over the CGaMP injection site and secured with dental cement. Following 3 weeks of viral expression, GCaMP fluorescence was detected through the optic fiber using a fiber photometry system (Neurophotometrics), with all data recorded in Bonsai visual reactive programming language (Lopes et al., 2015). Photobeam breaks detected with the drinking spout were aligned to the fiber photometry recording using an Arduino Uno as a digital acquisition device (constructed in house), and mice were video-tracked throughout the experiment using Bonsai. Mouse speed was calculated from x y position in NeuroExplorer, and GCaMP signal was aligned to behavioral events. Licking behavior was detected by photo beam breaks. Speed peaks were identified as local maxima in the calculated velocity trace that exceeded 5cm/s. We then scanned backwards in time from this peak in 0.1s bins to identify the first time-point above 2cm/s, which was considered the initiation of the locomotion bout. Locomotor arrests were defined as the 0.1s bin immediately preceding a minimum 2 s period of no change in the x or y position (Fobbs et al., 2020). Permutation analyses were run 2000 times with assignments for high or low trials shuffled before each permutation; the distribution of the resulting differences in lick behavior is shown. Three mice were excluded for optic fiber placements outside the VP.

### Behavioral Assays

#### Reward Consumption

Mice were individually placed in a 30cm x 20cm behavioral arena (Lab Products Rat One Cage 2100™) and given 60 min/day to consume a highly palatable liquid (Nesquik^®^, Nestlé, Switzerland). The liquid was presented in a liquid feeding tube (Bioserv) in the center of the arena. Before beginning baseline testing, mice were habituated to the behavioral arena for one day.

Baseline testing continued for 9-15 days to ensure habituation. The amount of time the mice spent drinking the liquid was recorded using infrared break-beam sensors connected to a microcontroller (Arduino, Turin, Italy). Following the baseline testing, the mice entered the experimental phase. The experimental phase had three session types: paired stimulation, scrambled stimulation, and no stimulation. In paired stimulation, the drinking behavior was paired with 20 Hz blue light (473 nm) stimulation for the duration of the time spent drinking. In scrambled stimulation, 1 sec of stimulation at 20 Hz was delivered randomly throughout the hour-long session. The no stimulation sessions were the same as the baseline testing; the mice had optic fibers attached but no stimulation was delivered during that session. The mice were tested for three days for each session type. The order of sessions was counterbalanced between the mice. The amount of time spent drinking and timestamps of each drinking bout were recorded.

#### Orofacial Taste Reactivity

Mice were placed individually in a behavioral arena where they had access to a 5% sucrose solution. Tongue protrusions were videotaped at 140 fps and results were manually scored. The first six lick bouts of each sessions was scored over 3 sessions / mouse. Alternating interactions with the sucrose solution were paired with 20 Hz blue light stimulation (473nm), counterbalanced across mice.

#### Real Time Place Preference

Mice were individually run in a behavioral arena that had two established zones distinguished by wall patterns. The RTPP was run in three parts: a pretest session, a stimulation session, and a test session. In each session, the mice were attached to optic fibers and placed in the arena for 45 minutes. In the pretest session, the mice freely explored the two zones without any stimulation. In the stimulation session, mice would receive 20 Hz stimulation while in the “active” experimental zone, while there was no stimulation delivered in the other “inactive” zone. Active and inactive zones were counterbalanced across subjects. The amount of time mice spent in each of the zones was recorded with a USB-powered webcam and zones were detected using Bonsai programing software (Lopes et al., 2015).

#### Optogenetic ICSS Assay

Mice were acclimated to behavioral testing chambers equipped with a device composed of two nose pokes, adapted from FED (Nguyen et al., 2016, 2017). Pokes on the active port resulted in a tone and delivery of 2 seconds of 20 Hz stimulation through the optic fibers, pokes on the inactive nose poke had no effect; the position of the active nosepoke was counterbalanced between subjects. Mice completed 5 daily 60 minute sessions over 5 consecutive days.

#### Two-Bottle Choice Assay

Mice were acclimated to an operant chamber equipped with two identical bottles of chocolate milk during their dark cycle (Godynyuk et al., 2019). Following this habituation, two identical bottles were again presented but licks registered on one bottle triggered optogenetic stimulation at 20 Hz, 4 ms pulses for the duration of the lick bout, with a minimum duration of 1 sec.

#### Open Field Test

Mice were individually run in a behavioral arena (50 cm x 50 cm x 30 cm). Mice were placed in the center of the arena and locomotor activity was recorded for 20 minutes. Afterwards, mice received 20Hz blue light stimulation (473 nm) for 20 minutes while locomotion was recorded. The behavior was recorded with a USB-powered webcam while distance traveled was detected using Bonsai.

### Data Analysis

Patch clamp electrophysiology data was analyzed offline in Clampex version 11. In vivo electrophysiological and fiber photometry data were analyzed in NeuroExplorer, to align events to neural activity data. Statistics were performed in Python 3.7, or Graphpad Prism version 8. A p value of 0.05 was considered significant, statistical tests are indicated throughout.

---

**Supplemental Figure 1.**
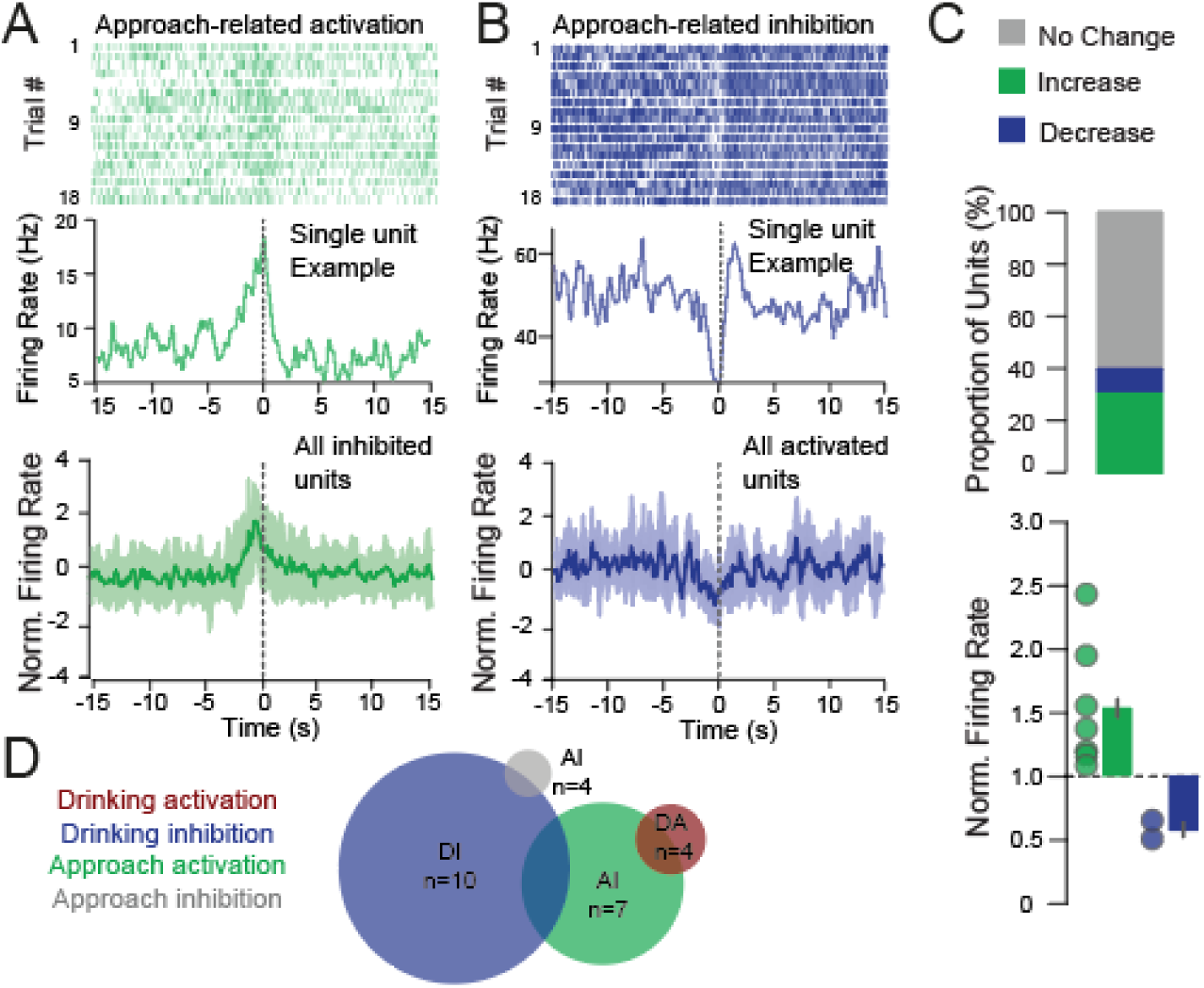
NAcSh units are activated by reward approach. **(A-B)** Top: examples of approach-inhibited and approach activated single units from the same mouse, aligned to drinking onset. Raster and PSTH of units inhibited (C) and activated (D) during reward approach; bottom = mean ± SEM. **(C)** Summary of approach-related responses of NAcSh units; 7.14% decrease, 25.0% increase, 67.86% no change, and normalized firing rate changes in response to approach (mean increase = 1.81 ± 0.20, mean decrease = 0.79 ± 0.09). **(E)** Euler diagram showing overlap of functionally defined NAcSh units.

**Supplemental Figure 2.**
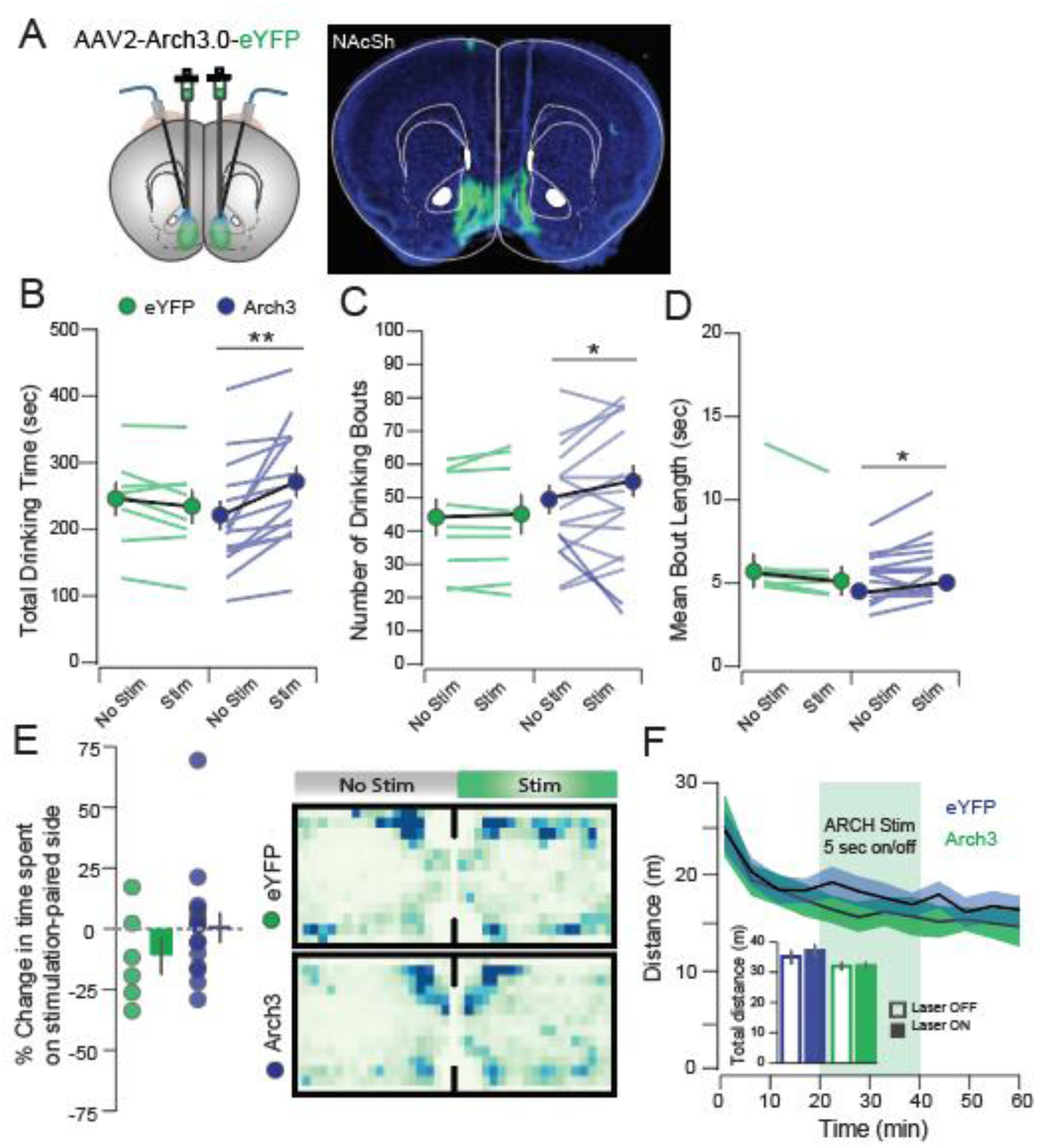
NAcSh inhibition promotes reward consumption without inducing a place preference or locomotor effects. **(A)** Histology showing viral infection and optic fiber placement. **(B)** Arch stimulation increased total drinking time (Arch No Stim: 217.93±21.57 sec, Arch Stim: 267.95±22.93 sec, t_14_ = 3.77, p = 0.002; eYFP No Stim: 24.422±30.03 sec, eYFP Stim: 231.43±25.55 sec, t_7_ =1.68, p=0.14), **(C)** number of bouts (Arch No Stim: 48.88±4.29, Arch Stim: 53.63±4.69 sec, t_14_ = 2.47, p = 0.038; eYFP No Stim: 43.21±5.40, eYFP Stim: 44.13±6.00, t_7_ =0.88, p=0.41) and **(D)** mean bout length (Arch No Stim: 4.63 ± 0.37 sec, Arch Stim: 5.16±0.46 sec, t_14_ = 2.27, p = 0.040; eYFP No Stim: 5.836 ± 0.847 sec, eYFP Stim:5.836±0.847, t_7_ =2.28, p = 0.06). **(E)** Arch3 inhibition did not induce a place preference in an RTPP task (eYFP: −11.92±7.71%, Arch: 1.06±5.96, t_19_=1.22, p=0.24) or **(F)** alter locomotor activity in an open field task (eYFP No Stim: 320.97±13.41 m, eYFP Stim: 323.41±12.02 m, Arch No Stim: 353.12±22.46 m, Arch Stim: 374.07±21.34 m, F_StimxVirus;1,19_=1.33, p=0.26). *p<0.05, **p < 0.01

**Figure S3.**
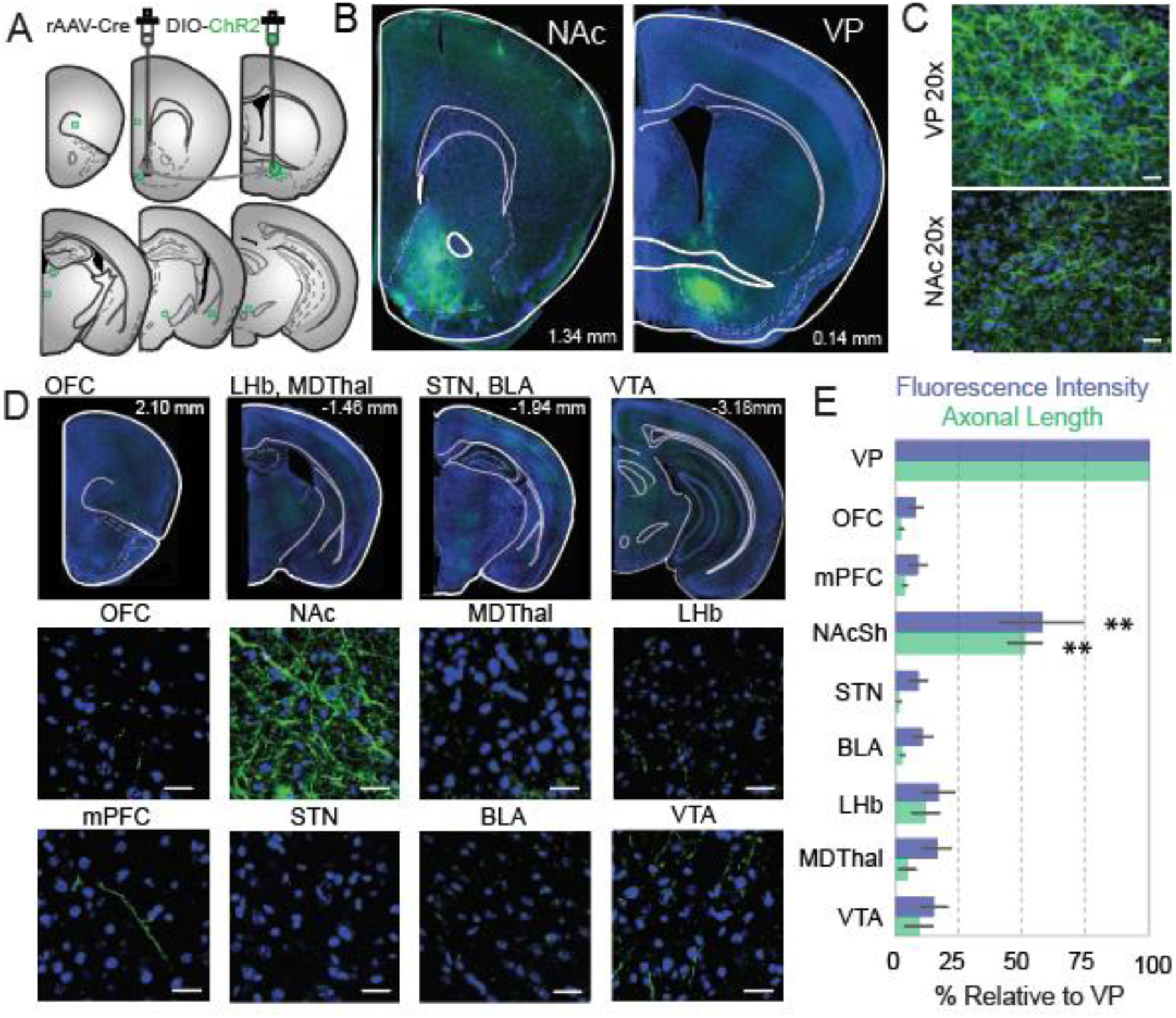
Retrograde viral tracing reveals vArky fibers preferentially in the NAcSh. **(A)** rAAV-Cre was injected in the NAcSh, DIO-ChR2 was injected into the VP and serial sections were taken of known VP projection sites. **(B-C)** 5x and 20x overview of ChR-labeled terminals in the NAcSh and infected cell bodies in the VP. **(D)** 10x representative images of serial sections (top) and 20x confocal images of canonical VP projection areas. **(E)** Quantification of fluorescence density (eYFP) and total axonal length, normalized to VP (n=4 sections/brain region, 6 mice). Intensity and axonal length in the NAcSh were significantly greater than any area examined. Abbreviations: Nucleus accumbens shell (NAcSh), ventral pallidum (VP), medial prefrontal cortex (mPFC), orbitofrontal cortex (OFC), basolateral amygdala (BLA) lateral habenula (LHb), mediodorsal thalamus (MDThal), subthalamic nucleus (STN), ventral tegmental area (VTA). F_length_=148.4 p<0.001, F_fluorescence_=112.6. Scale bars=25 µm.

**Figure S4.**
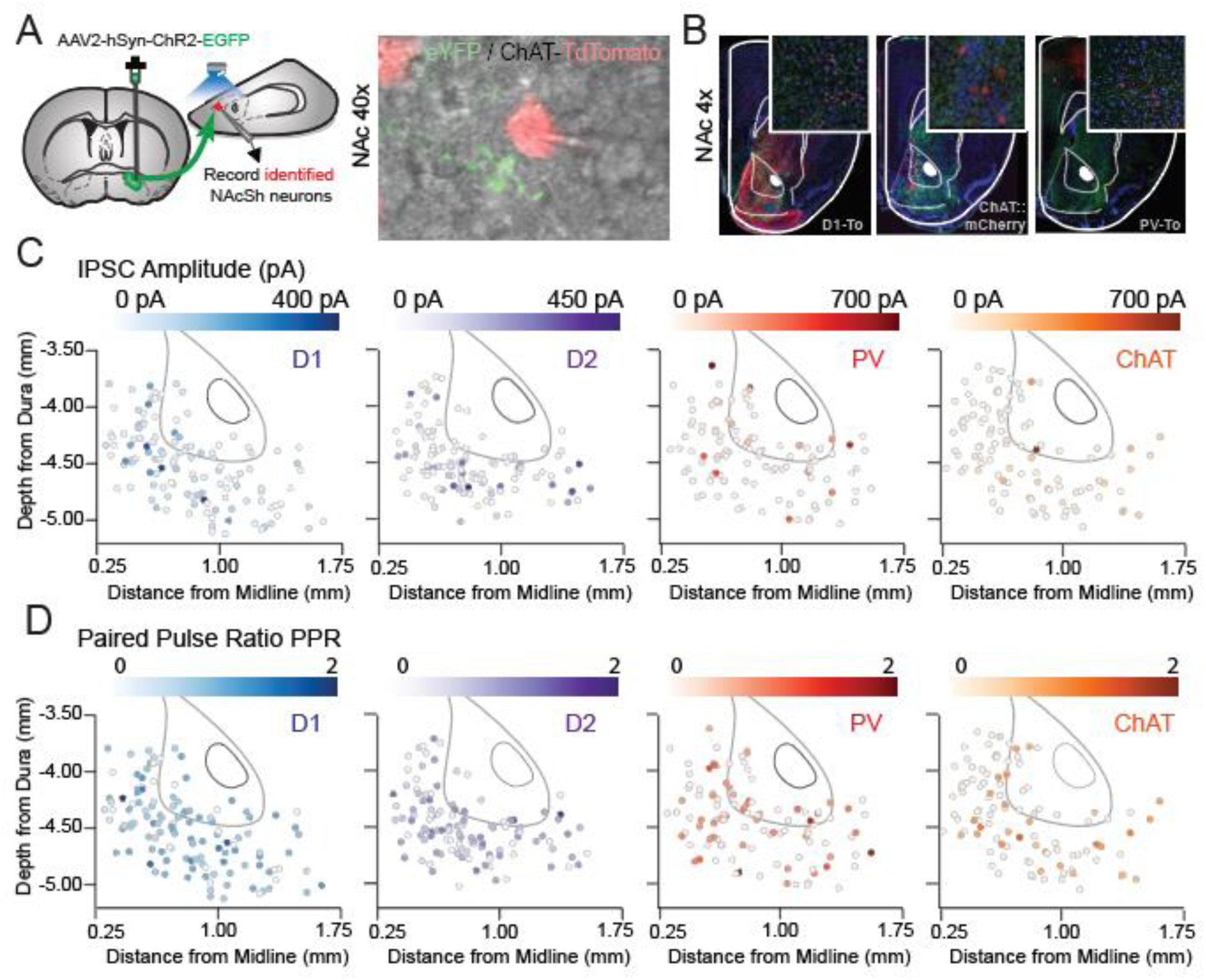
Topography of release properties of VP inputs to NAcSh. **(A)** Merged fluorescent image of recorded tomato-expressing neuron adjacent to EYFP-labeled terminals from the VP during patch-clamp experiments. **(B)** 4x image of ChR2-injection site in VP and terminal fields in the NAc in each of three reporter lines. **(C-D)** Amplitude **(C)** and PPR **(D)** of oIPSC expressed as location of recorded neuron within the NAcSh in D1-MSNs (n=118 cells / 11 mice), D2-MSNs (n=117 cells / 12 mice), PVs (n=112 cells / 12 mice), and CINs (n=104 cells / 10 mice).

**Figure S5.**
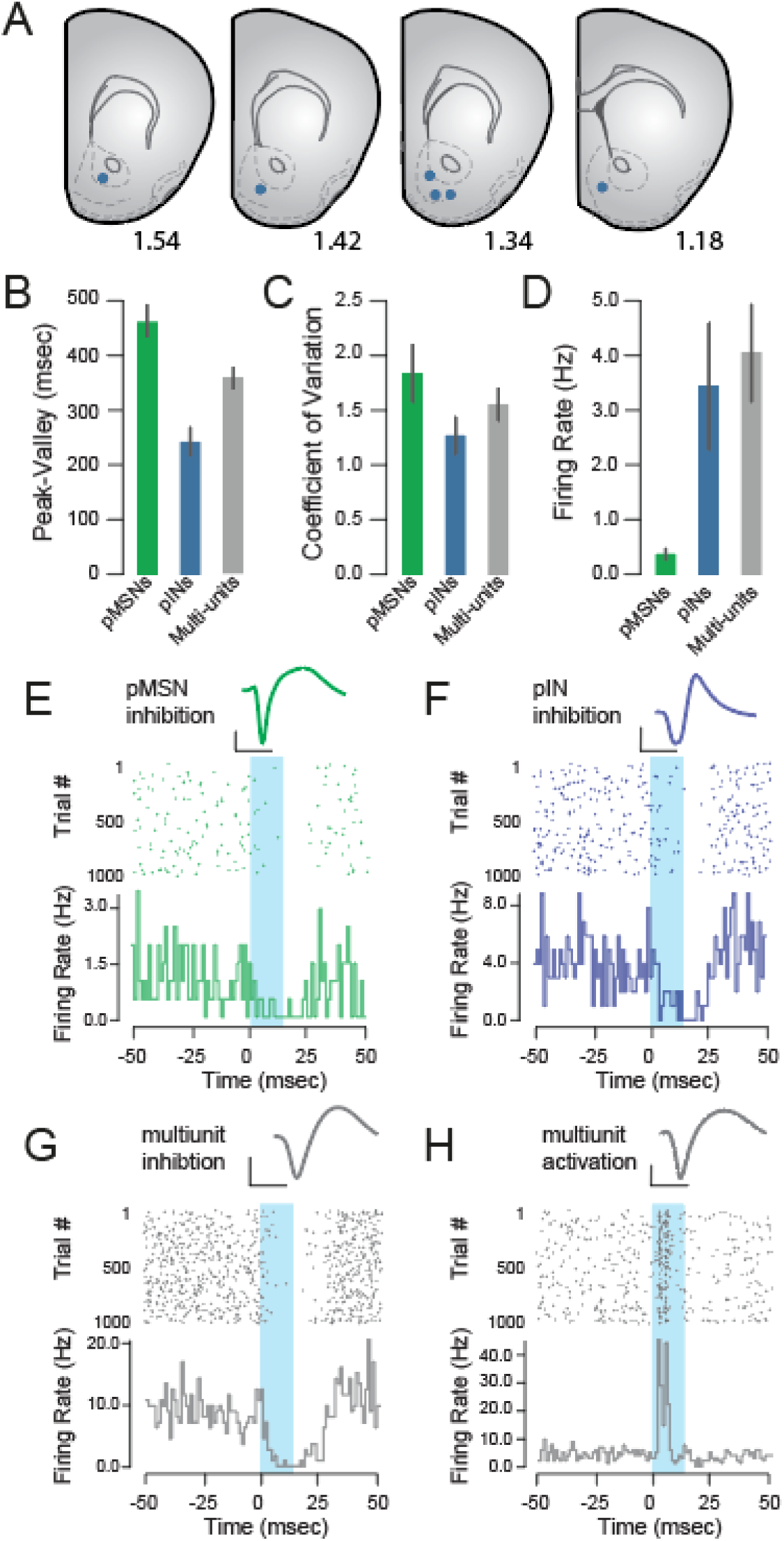
Properties of NAcSh neurons and in vivo responses to activation of vArky terminals. **(A) Location of recording arrays in the NAcSh. (B-D)** Units were classified according to peak-valley width (pMSN: 462.8±27.96 µs, pIN: 242.01±26.27 µs), CV (pMSN: 1.77±0.254, pIN: 1.23±0.17) and firing rate (pMSN: 0.40±0.11 Hz, pIN: 3.53±1.20 Hz). **(E-H)** representative waveform and PSTH of single neural examples in response to 15 ms of vArky stimulation. Scale bar = 500µs, 50µV.

**Figure S6.**
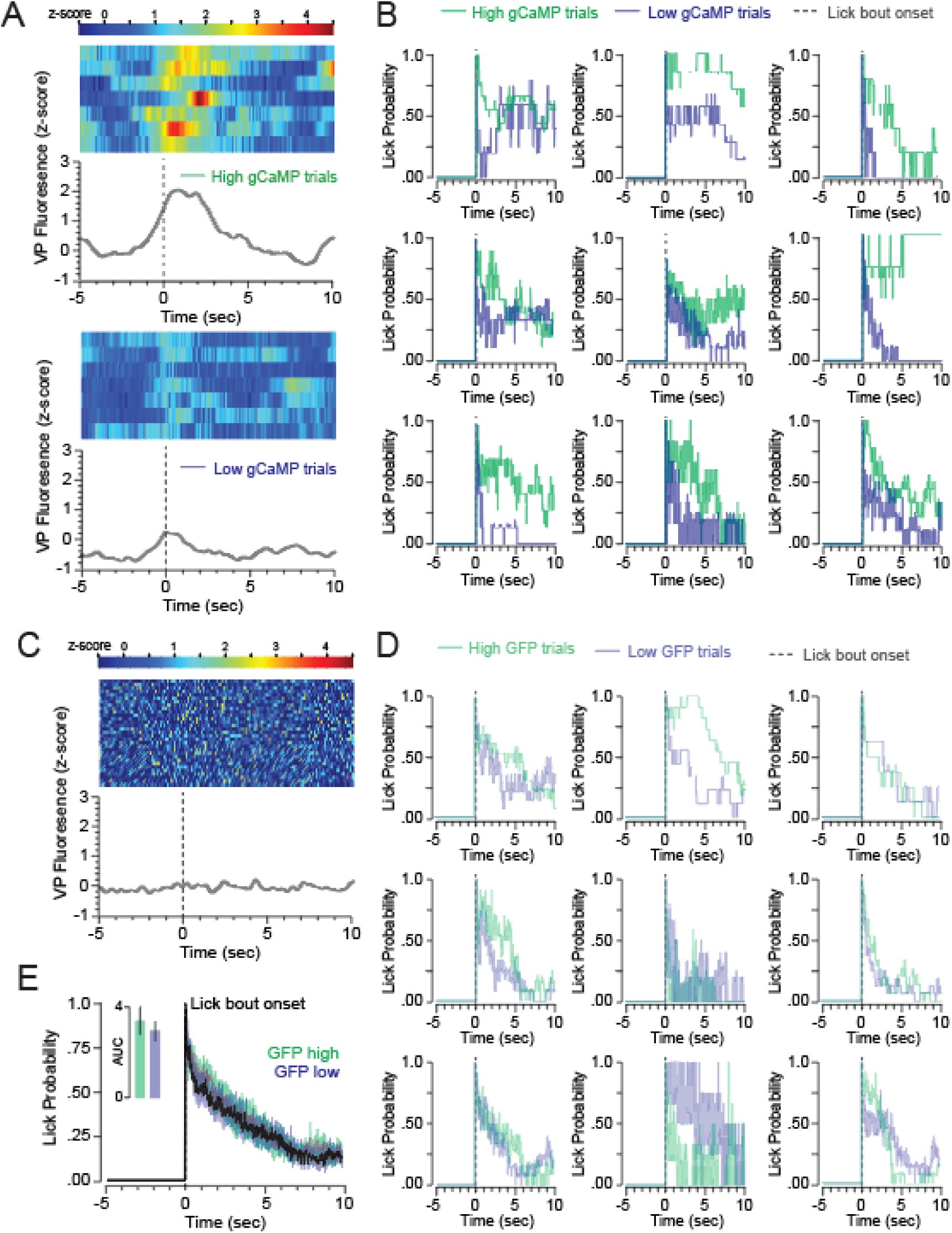
Lick behavior of individual subjects, as a function of high- vs. low-VP calcium signals. **(A)** Single-subject heat map of arkypallidal fluorescence signal across high GCaMP (top) and low GCaMP (bottom) trials aligned to reward consumption onset. **(B)** Lick behavior of high-vs. low-VP calcium trials of all individual GCaMP-injected subjects. **(C)** Single-subject heat map of fluorescence signal of GFP mouse. **(D)** Lick behavior of high-vs. low-VP signal trials of all GFP-injected subjects, **(E)** Average lick behavior of high and low VP signal trials of all GFP-injected mice (AUC lick probability high signal trials: 3.16±0.65, AUC lick probability low signal trials: 2.94±0.49).

**Figure S7.**
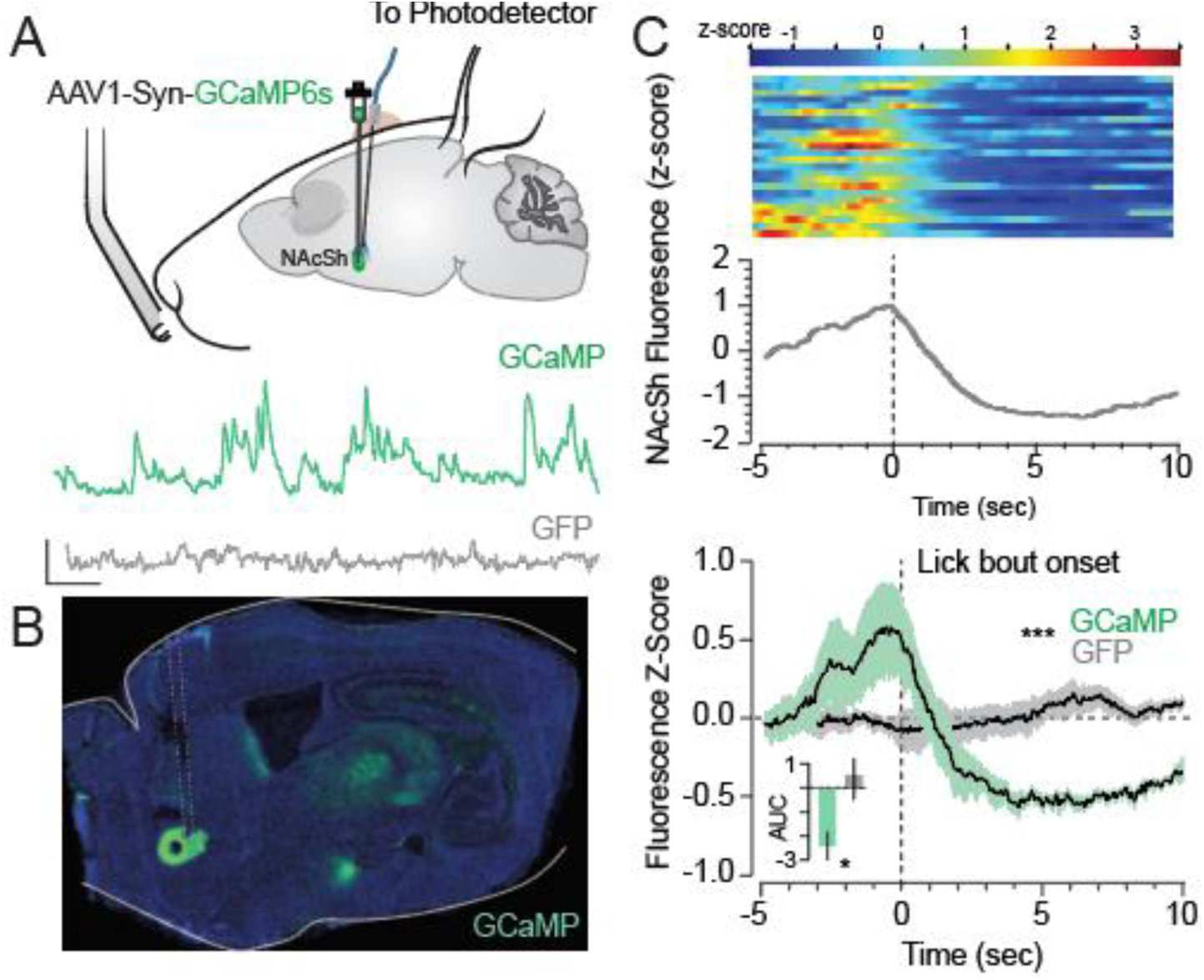
Calcium activity in the NAcSh decreases upon reward consumption onset. **(A-B)** GCaMP injection in the NAcSh, representative traces (scale bar: 10 sec, 1 z-score) and histology. **(C)** Single-subject heat map of NAcSh GCaMP signal and **(D)** combined signal from all subjects (n=6/6 GCaMP/GFP) aligned to reward consumption onset (mean AUC GCaMP: −2.58±0.65, GFP: −0.14±1.40, t_11_=2.87, p=0.015).

**Figure S8.**
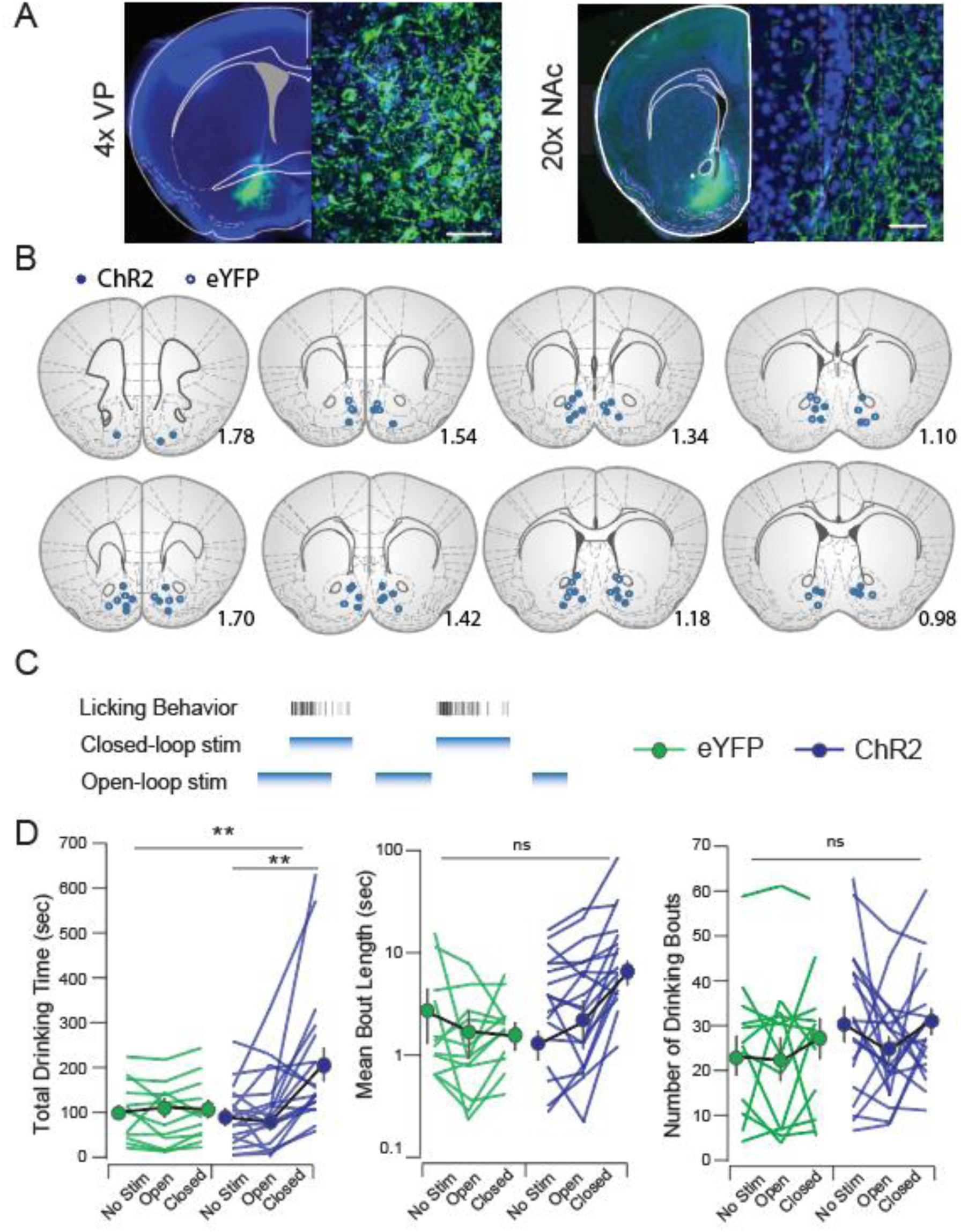
Histological verification and behavior control with open-loop optogenetic stimulation. **(A)** Infection site in VP and terminal fields in NAcSh; scale bar=50µM. **(B)** Verified placements of optic fibers **(C)** Schematic of open-vs. closed-loop stimulation **(D)** Individual subjects and mean±SEM of total drinking time (eYFP_NoStim_: 85.98±20.47, eYFP_Open_: 85.98±12.46, eYFP_Closed_: 91.10±19.03, ChR2_NoStim_: 91.50±15.58, ChR2_Open_: 101.56±13.76, ChR2c_losed_: 195.81±38.51, F_virus•stim_=4.47, p=0.016), bout length (eYFP_NoStim_: 5.34±1.87, eYFP_Open_: 5.34±1.87, eYFP_Closed_ = 4.76 ± 1.16, F_virus•stim_=3.73, p=0.030) and number of bouts (eYFP_NoStim_: 22.86±4.57, eYFP_Open_: 21.85±4.97, eYFP_Closed_: 22.86±4.78, ChR2_NoStim_: 30.17±4.02, ChR2_Open_: 24.26±2.88, ChR2_Closed_: 30.21±2.89, F_virus•stim_=0.263, p=0.77). *p<0.05, **p<0.01.

